# A chemically inducible multimerization system for tunable and background-free RTK activation

**DOI:** 10.1101/2025.09.06.674232

**Authors:** Yuanmin Zheng, Jinyu Fei, Abhirup Chakrabarti, Ruobo Zhou

**Affiliations:** Department of Chemistry, The Pennsylvania State University, University Park, PA 16802, USA; Department of Biomedical Engineering, The Pennsylvania State University, University Park, PA 16802, USA; Department of Biochemistry and Molecular Biology, The Pennsylvania State University, University Park, PA 16802, USA; The Huck Institutes of Life Sciences, The Pennsylvania State University, University Park, PA 16802, USA

## Abstract

Receptor tyrosine kinases (RTKs) are key regulators of diverse cellular processes, including differentiation, migration, proliferation, survival, and intracellular communications, by transducing extracellular cues into intracellular responses. Upon oligomerization at the plasma membrane, RTKs become activated and initiate major downstream signaling cascades, such as the ERK pathway, which modulates cytoskeletal dynamics through phosphorylation of cytoskeletal regulators, regulation of actin-binding proteins, and transcriptional activation of immediate-early genes involved in cell structure and motility. Light-inducible RTK systems have been developed to achieve spatiotemporal control of RTK clustering and activation for both basic cell biology research, and engineered applications, such as controlling cell migration, proliferation, or differentiation. However, these systems are limited by high basal RTK activation, where substantial RTK activation occurs even before induction, leading to unintended ERK activation and downstream effects. Here, we report a chemically inducible RTK platform that minimizes basal activation while enabling direct visualization of RTK clustering at the plasma membrane upon induction. Single-cell imaging reveals visible RTK clusters after induction, with total RTK abundance in the clusters correlating with ERK phosphorylation levels. Using this system, we achieved precise and rapid control over multiple ERK-dependent cellular processes, including disassembly of spectrin-based membrane skeleton and nuclear entry of transcription factors STAT3 and CREB, while maintaining minimal basal activity before induction. In contrast to previously developed inducible RTK systems, which can perturb cytoskeletal structures or transcription factor dynamics even without stimulation, our design preserves native cellular architecture and nuclear signaling until activation is intentionally triggered. Collectively, these results establish our system as a robust and versatile platform for dissecting RTK signaling dynamics and engineering cell behaviors with precise, on-demand spatiotemporal control.

## INTRODUCTION

Receptor tyrosine kinases (RTKs) comprise a structurally diverse family of transmembrane proteins that function as key regulators of cellular signaling, by converting extracellular stimuli into coordinated intracellular responses^1–4^. Ligand binding to RTKs induces conformational rearrangements that facilitate dimerization or higher-order multimerization of their cytoplasmic kinase domains, leading to autophosphorylation of specific tyrosine residues and subsequent recruitment of downstream signaling effectors^5,6^. Among the well-characterized downstream pathways of RTKs is the mitogen-activated protein kinase (MAPK) cascade, particularly the Raf–MEK–ERK axis, which governs essential cellular processes including proliferation, differentiation, morphogenesis, and cytoskeletal remodeling^7–9^. Notably, the amplitude and duration of ERK activation are highly sensitive to the spatial organization and clustering state of RTKs at the plasma membrane^10,11^. Parameters such as receptor density, oligomerization state and geometry, and subcellular localization may contribute to modulating downstream signaling strength in highly context-dependent manners^12,13^.

Despite growing interest in controlling RTK clustering to modulate downstream cellular effects, such as controlling cell migration, proliferation, or differentiation, current tools for manipulating receptor multimerization remain limited in precision^14,15^. Optogenetic systems, particularly those based on light-inducible oligomerization domains CRY2^16^ or iLID^17^, have been widely employed to induce RTK clustering in a light-dependent manner. However, these platforms often exhibit substantial basal activation even in the absence of light illumination^18^, which is further exacerbated at high expression levels, owing to the intrinsic tendency of light-inducible oligomerization domains to self-associate. This leaky activation complicates quantitative analyses of downstream signaling, such as resolving dose–response relationships or defining kinetic thresholds with high confidence. Moreover, optogenetic activation requires continuous or repeated light illumination, which can introduce phototoxic effects and prove impractical in certain *in vivo* contexts. Chemically induced dimerization (CID) systems, such as rapamycin-mediated pairing of FK506-binding protein (FKBP) and the FKBP–rapamycin binding (FRB) domain of mTOR^19,20^, offer a light-independent alternative for triggering RTK activation. However, conventional CID approaches are generally restricted to dimerization and do not support tunable higher-order assemblies, thereby limiting control over clustering geometry, valency, and spatial localization. These limitations underscore the need for a chemically controlled, modular system that enables background-free activation, and precise tunability, enabling detailed investigation of how RTK spatial organization translates into analog outputs such as ERK phosphorylation.

To overcome the limitations of existing RTK clustering tools, we developed a chemically inducible platform for precise control of RTK multimerization with minimal background activity, using fibroblast growth factor receptor 1 (FGFR1)^21–24^ as a prototypical model receptor. Our system leverages rapamycin-mediated heterodimerization between FKBP and FRB, a well-characterized CID pair that functions independently of light and exhibits rapid dimerization kinetics²⁷. To achieve higher-order receptor clustering beyond simple dimerization, we integrated engineered oligomerization tags HOTag3 and HOTag6, two previously reported coiled-coil domains that assemble into hexametric and tetrameric complex, respectively^25,26^. By fusing the intracellular domain of FGFR1 to various combinations of FKBP/FRB and HOTag3/HOTag6 modules, we generated a library of chimeric receptor constructs with defined clustering geometries and subcellular localizations. Systematic evaluation revealed that configurations with membrane-anchored FGFR1-FKBP-HOTag6 paired with cytosolic FRB-HOTag3 supported strong rapamycin-inducible clustering while maintaining negligible basal ERK activation. For benchmarking, we compared our optimized constructs to reported CRY2-based optogenetic systems. Optogenetic FGFR1 systems based on the CRY2 variants—including wild type(WT)CRY2, CRY2olig^27^ and optodroplet-forming CRY2 fusions^28^—exhibited considerable basal ERK activation even without light stimulation, consistent with the previously reported properties that CRY2 may exhibit some degree of self-association in the dark, even without light stimulation^29^. In contrast, our rapamycin-inducible system showed minimal basal ERK phosphorylation and robust, tunable activation upon rapamycin treatment. Live-cell imaging further confirmed that rapamycin rapidly induced spatially localized FGFR1 clustering, accompanied by recruitment of cytosolic components to the plasma membrane^30–33^. These optimized constructs enable on-demand, higher-order FGFR1 assembly that robustly triggers downstream ERK signaling. Importantly, the modular architecture of this system allows independent tuning of clustering localization, providing a versatile platform with significant advantages over existing optogenetic inducible approaches.

Using this modular chemical platform, we systematically investigated how the degree of FGFR1 clustering regulates ERK signaling in human cells. FGFR1 serves as an effective model system for studying the biophysical determinants of signal activation, due to its relevance in development and disease, and its pronounced sensitivity to multimerization^21–24^. The quantitative relationship between FGFR1 clustering degree and the resulting strength of ERK signaling remains unclear, posing an important challenge for both mechanistic studies of cell signaling and the rational engineering of synthetic signaling systems^21,24,34,35^.Our quantitative single-cell analysis showed that the extent of FGFR1 multimerization, measured by the total amount of FGFR1 molecules within FGFR1 clusters per cell, strongly correlated with phosphorylated ERK levels, establishing a proportional relationship between receptor clustering and downstream signaling amplitude.

To demonstrate the advantages of our optimized chemical-inducible RTK system, we utilized it to control multiple ERK-dependent cellular outputs. Previous studies have shown that following ERK activation, the spectrin-based membrane skeleton^36,37^ can be rapidly degraded^38,39^ and that transcription factors, such as STAT3 and CREB, enter the nuclear to control transcription activites. Using our system, we showed that rapamycin-induced FGFR1 clustering led to membrane skeleton disassembly and significantly increased nuclear entry of STAT3 and CREB in cells expressing our chemical-inducible FGFR1 constructs, whereas untreated cells retained intact membrane skeleton structures and minimal nuclear entry of STAT3/CREB. In contrast, cells expressing CRY2-based optogenetic systems exhibited membrane skeleton disassembly and enhanced nuclear entry of STAT3/CREB even without any light stimulation. Collectively, our results underscore our system’s capacity for precise, on-demand inducible control of RTK clustering, enabling interrogation of both cytoplasmic and nuclear ERK-dependent outputs while maintaining exceptionally low basal activity.

## MATERIALS AND METHOD

### Molecular cloning and plasmid construction

All plasmids and DNA fragments used in this study were obtained from Addgene or synthesized by commercial providers. Constructs were assembled using Gibson assembly (New England Biolabs, E5510S) according to the manufacturer’s instructions.

The coding sequence for human FGFR1 was obtained from Addgene plasmid #116740. CRY2PHR (CRY2) and its oligomerization-prone mutant CRY2olig were derived from Addgene plasmid #106169. The Optodroplet system was constructed using Addgene plasmid #111507 as a backbone, in which the original FusionRed fluorophore was replaced with EGFP via PCR amplification and Gibson assembly. Cytoplasmic FGFR1 was fused at its C-terminus to CRY2, CRY2olig, or Optodroplet to generate the FGFR1–CRY2, FGFR1–CRY2olig, and FGFR1–Optodroplet constructs, respectively.

For the rapamycin-inducible clustering system, coding sequences for FKBP and FRB were obtained from Addgene plasmid #40896. The HOTag3 and HOTag6 multimerization domains were sourced from Addgene plasmids #106924 (FKBP–EGFP–HOTag3) and #106919 (FRB–EGFP–HOTag6), respectively. EGFP was replaced with mScarlet in both constructs by Gibson assembly to yield FKBP–mScarlet–HOTag3 and FRB–mScarlet–HOTag6. Additional variants incorporating alternative domain arrangements and linker lengths were assembled to generate constructs (Pairs 1 through 10) for systematic evaluation of FGFR1 clustering efficiency.

### Cell culture and transfection

Human bone osteosarcoma epithelial cells (U2OS; ATCC HTB-96) were cultured in Dulbecco’s Modified Eagle Medium (DMEM; Corning, 10-013-CV) supplemented with 10% (v/v) fetal bovine serum (FBS; Gibco, A52567-01) and 1% (v/v) penicillin–streptomycin (Gibco, 10378-016). Cells were maintained at 37 °C in a humidified incubator with 5% CO₂. For passaging, cultures were dissociated using 0.25% Trypsin–EDTA (Corning, 25-053-CI). For imaging experiments, cells were seeded onto 18-mm round glass coverslips (Electron Microscopy Sciences, 72222-01) placed in 12-well tissue culture plates.

Plasmid transfection was performed using TransIT®-LT1 reagent (Mirus Bio, MIR2300) according to the manufacturer’s protocol. At 60–80% confluency, cells were transfected with 1.0 µg of plasmid DNA diluted in 100 µL of Opti-MEM (Gibco, 31985-070), followed by addition of 3 µL of TransIT-LT1 reagent. The mixture was incubated at room temperature for 15 min to allow complex formation, then added dropwise to the culture medium. After 6 h, the medium was replaced with fresh DMEM supplemented with 10% FBS to promote recovery and transgene expression. Cells were either imaged live or fixed for downstream analysis 24-48 h post-transfection, depending on the experimental protocol.

### Blue light stimulation

For optogenetic clustering assays, U2OS cells expressing CRY2-based constructs were stimulated using a custom-built blue light system, as previously reported1. The setup consisted of a standard blue LED light source (465 nm) powered by a DC power supply. LED brightness was controlled to the desired level. Cells were stimulated with continuous blue light exposure for defined durations ranging from 30 seconds to 5 minutes, depending on the experimental conditions. To prevent unintended activation, samples were protected from ambient blue light prior to stimulation.

Live-cell imaging during stimulation was performed using Gibco™ Live Cell Imaging Solution (Thermo Fisher, A14291DJ), and cells were maintained at 37 °C throughout the assay. For fixed-cell experiments, light stimulation was followed by immediate fixation with 4% (w/v) paraformaldehyde (PFA) containing 4% (w/v) sucrose in phosphate-buffered saline (PBS) prior to immunostaining.

### Rapamycin stimulation

Rapamycin (Millipore Sigma, Cat. No. 553210) was reconstituted in DMSO to a stock concentration of 100 mM, aliquoted, and stored at –20 °C. For live-cell imaging, 100 mM rapamycin stock was diluted to a final working concentration of 500 nM in Gibco™ Live Cell Imaging Solution (Thermo Fisher Scientific, Cat. No. A14291DJ). U2OS cells were pre-equilibrated by rinsing with pre-warmed imaging solution and incubating at 37 °C for at least 15 minutes prior to stimulation. Rapamycin was then added directly to the imaging chamber during acquisition, and time-lapse confocal imaging was performed over a 10-30 minute window post-treatment to monitor clustering dynamics. For fixed-cell imaging, cells were treated with 500 nM rapamycin in DMEM containing 10% FBS and 1% penicillin–streptomycin for the indicated durations, after which they were fixed and immunostained.

### Cell fixation and immunostaining

U2OS cells grown on glass coverslips were fixed with 4% (w/v) PFA containing 4% (w/v) sucrose in PBS for 20 minutes at room temperature. After fixation, cells were rinsed three times with PBS and permeabilized with 0.2% (v/v) Triton X-100 in PBS for 10 minutes. Cells were then blocked with 3% (w/v) bovine serum albumin (BSA) in PBS for 1 hour at RT. Primary antibodies were diluted in blocking buffer and incubated with samples overnight at 4°C. The following day, cells were washed with PBS three times and incubated with fluorophore-conjugated secondary antibodies in blocking buffer for 1 hour at RT. After final washes in PBS, coverslips were mounted for imaging.

For immunostaining of phosphorylated ERK (p-ERK), U2OS cells were fixed at designated time points following light or rapamycin stimulation. Cells were first treated with 4% (w/v) PFA containing 4% (w/v) sucrose in PBS for 20 minutes at RT, followed by ice-cold methanol incubation at −20°C for 10 minutes. Samples were subsequently permeabilized with 0.2% Triton X-100 in PBS for 10 minutes and then subjected to blocking and antibody staining as described above.

The following primary antibodies were used in this study: mouse anti-βII spectrin antibody 1:200 dilution for immunofluorescence (Santa Cruz Biotechnology, sc-136074 or sc-515592); rabbit anti-phospho-ERK1/2 (Thr202/Tyr204) antibody 1:300 dilution for immunofluorescence (Cell Signaling Technology, 4370S). The following secondary antibodies were used in this study: Alexa Fluor-647-conjugated donkey anti-mouse IgG antibody 1:800 for IF (Jackson ImmunoResearch, 715-605-151); Alexa Fluor-647-conjugated donkey anti-rabbit IgG antibody 1:800 for IF (Jackson ImmunoResearch, 711-605-152).

### Confocal imaging

Confocal imaging was performed on a custom microscope built on an Olympus IX-83 inverted body, equipped with a CrestOptics X-Light V3 spinning disk confocal unit. A 60× (NA 1.42) UPlanXApo or 100× (NA 1.40) UPlanSApo oil immersion objective (Olympus) was used in combination with an ORCA-Fusion BT digital CMOS camera (Hamamatsu, C15440-20UP). Illumination was provided by an LDI-NIR-7 Laser Diode Illuminator (89 North), enabling sequential excitation of fluorophores including AF488 (or EGFP), mScarlet, and AF647.

For each field of view, *z*-stack images were acquired across the full cellular volume, with a step size of 0.3-0.5 μm. To generate the images shown in the main figures, all *z*-stacks were processed using maximum-intensity projection. Projection was performed in MATLAB using the built-in max() operation applied along the *z*-dimension, yielding a 2D representation of the maximum-intensity values throughout the cellular volume.

### 3D-STORM imaging

Single-color three-dimensional stochastic optical reconstruction microscopy (3D-STORM) was performed using a custom-built setup based on a Nikon Eclipse-Ti2 inverted microscope, equipped with a 100× oil-immersion objective (NA 1.45, Plan Apo, Olympus) and an electron-multiplying CCD camera (Andor iXon Life 897). A 642-nm laser (MPB Communications Inc., 2RU-VFL-P-2000-642-B1R; 2000 mW) and 405-nm (Coherent, OBIS 405 nm LX, 1265577; 140 mW) was introduced through the rear port of the microscope and directed toward the edge of the objective to achieve near-total internal reflection (TIR) illumination, enabling selective excitation of fluorophores within a few microns of the coverslip surface.

For STORM imaging, AF647 was used as the reporter fluorophore. A cylindrical lens was inserted into the detection path to introduce point spread function ellipticity, allowing axial (*z*-axis) localization of single molecules based on the shape of their emission profile^40^.

Cells were imaged in a oxygen-scavenging STORM imaging buffer composed of 100 mM Tris-HCl (pH 7.5), 100 mM cysteamine (Sigma-Aldrich), 5% (w/v) glucose, 0.8 mg/mL glucose oxidase (Sigma-Aldrich), and 40 μg/mL catalase (Sigma-Aldrich). AF647 molecules were continuously excited with the 642-nm laser (∼2 kW/cm²), switching them into a non-fluorescent dark state. A low-intensity 405-nm laser was used for controlled reactivation to maintain a sparse population of active fluorophores suitable for single-molecule localization.

A typical 3D STORM image was reconstructed from ∼30,000 frames acquired at 110 Hz. Image reconstruction and molecule localization were performed using previously established algorithms2, rendering each localization as a 2D Gaussian peak centered at the estimated molecular position. Final images were visualized using custom MATLAB or ImageJ-based scripts.

### Data quantification and analysis

All subsequent data quantification and analyses were performed using custom MATLAB scripts. To quantify the intensity of p-ERK and spectrin in transfected (EGFP-positive) cells, we first generated binary cell masks from the EGFP channel. The EGFP signal was smoothed using a Gaussian filter (σ = 2), followed by adaptive thresholding to generate binary masks. Hole filling was applied on a per-connected-component basis to ensure enclosed cellular regions were captured. The resulting masks were further refined by removing small objects (<5000 pixels) and morphological closing. Individual cells were labeled, and corresponding p-ERK and spectrin images were loaded from the same field of view (FOV). For each labeled EGFP-positive cell, the mean and standard deviation (SD) of pixel fluorescence intensities were calculated within the segmented cell mask in each respective channel across the FOV. The coefficient of variation (CV) was then computed as the ratio of SD to the mean fluorescence intensity, providing a measure of signal heterogeneity. In parallel, EGFP puncta were detected using a top-hat filter (disk size = 10 pixels) followed by a 99th percentile threshold and size filtering to exclude small artifacts (<3 pixels). Puncta count, mean and SD values, and CVs for p-ERK and spectrin intensities in each cell were compiled and exported into Excel for downstream analysis.

### Statistical Analysis

All statistical analyses were performed using MATLAB. Unless otherwise noted, all experiments were conducted with three biological replicates. For each replicate, 30–50 EGFP-positive cells were imaged and quantified. Data are reported as mean ± s.d..

For measurements of p-ERK, βII-spectrin, p-STAT3, and p-CREB, each biological replicate produced one averaged value calculated from 30–50 cells, and therefore each data point represents one biological replicate (*n* = 3). Background correction was performed using the mean signal of neighboring untransfected cells within the same field of view prior to averaging. Statistical significance for these replicate-level comparisons was determined using unpaired two-tailed Student’s t-tests.

For puncta and cluster-associated measurements, each data point represents a single cell. Values from all analyzed cells across three biological replicates were pooled for statistical comparison. Group differences for these single-cell datasets were assessed using two-sided Welch’s t-tests, which do not assume equal variance between conditions.

## RESULTS AND DISCUSSION

### CRY2-based optogenetic FGFR1 constructs exhibit basal leaky ERK activation

Previously reported optogenetic RTK constructs have enabled light-inducible activation of downstream signaling cascades, including the MAPK/ERK pathway, with high spatiotemporal resolution^16^. Most of these systems utilize the CRY2-based clustering approach, which induces RTK oligomerization at the plasma membrane in response to blue light, thereby activating ERK cascades^41,42^. To evaluate the performance of CRY2-medicated clustering in regulating FGFR1 signaling, we tested three optogenetic FGFR1 constructs **(Figure 1a)** with distinct clustering properties: 1) FGFR1-CRY2, in which cytoplasmic portion of FGFR1 (CytoFGFR1) is fused to wild-type CRY2; 2) FGFR1-CRY2oligo, incorporating the E490R^43^ point mutation to enhance light-induced oligomerization; 3) FGFR1-Optodroplet, where Optodroplet is a previously developed module combining wild-type CRY2 with a C-terminal tail and an intrinsically-disordered FUS domain to achieve light-induced formation of liquid condensates of the tagged protein via liquid-liquid phase separation^27,44^. All constructs were N-terminally tagged with a myristoylation (Myr) motif for membrane anchoring and fused to EGFP for fluorescence imaging.

**Figure 1.**
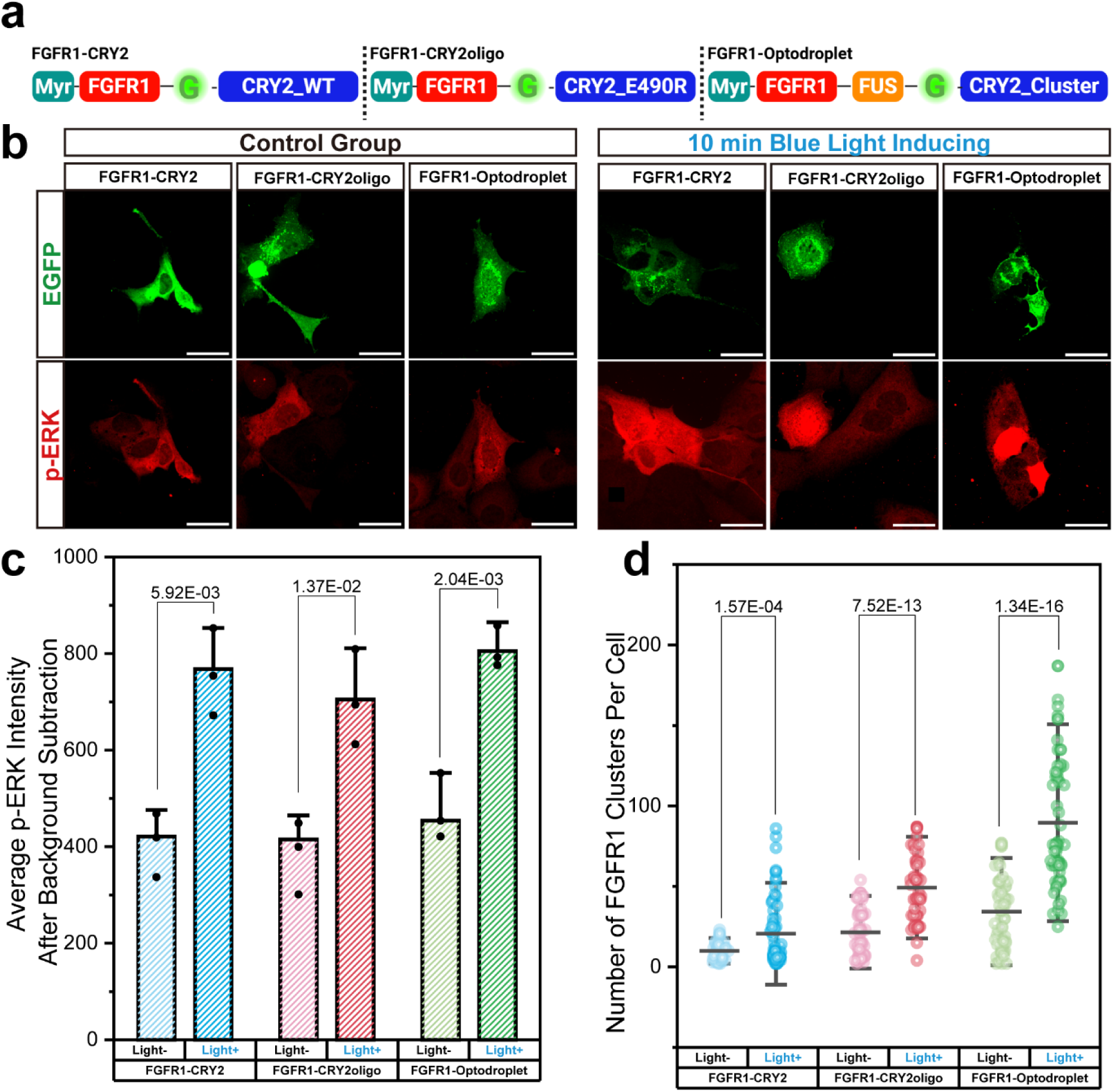
CRY2-based optogenetic clustering of FGFR1 activates ERK signaling. **(a)** Schematic diagrams of three optogenetic FGFR1 constructs. **(b)** Confocal immunofluorescence images of fixed U2OS cells expressing each construct. Scale bars, 10 μm. **(c)** Quantification of average total p-ERK fluorescence intensity in individual transfected cells with or without light stimulation. **(d)** Quantification of FGFR1 cluster number.

U2OS cells transfected with the plasmids encoding each optogenetic FGFR1construct were either exposed to blue light for 10 minutes or maintained in the dark, followed by phospho-ERK (p-ERK) quantification using a previously established immunofluorescence-based assay^45^. All three constructs—FGFR1–CRY2, FGFR1–CRY2olig, and FGFR1–Optodroplet—exhibited robust increases in p-ERK signal upon light stimulation, confirming that light-induced FGFR1 clustering activates the MAPK/ERK pathway **(Figure 1b, right panels)**. However, we also observed elevated p-ERK signal in transfected cells maintained in the dark, compared to neighboring untransfected cells **(Figure 1b, left panels)**. To more accurately quantify induced ERK activation, we calculated the total p-ERK fluorescence intensity at the single-cell level after background correction, where background was defined as the average total p-ERK fluorescence intensity in untransfected cells **(Figure 1c)**. Statistical analysis revealed significant increases in p-ERK upon light stimulation for all constructs. However, elevated p-ERK levels in the unstimulated (light-) conditions, compared to the background p-ERK fluorescence determined from untransfected cells, suggest substantial basal leaky ERK activation without light illumination. Moreover, we quantified the number of EGFP clusters per cell in the EGFP-channel to assess receptor multimerization. All three constructs showed a significant increase in FGFR1 cluster number per cell upon light stimulation **(Figure 1d)**.

We next examined the time course of blue-light–induced ERK activation for FGFR1-CRY2oligo and FGFR1-Optodroplet. In cells transfected with either construct, the average p-ERK signal increased exponentially and reached a plateau within 10–15 minutes of light illumination (**Figure S1a)**. Interestingly, we also observed that blue-light illumination alone induced a time-dependent increase in ERK phosphorylation in untransfected cells (**Figure S1b**), indicating that the undesired basal ERK activation in optogenetic systems may also arise from the light stimulation itself.

To further test whether the basal (“leaky”) ERK activation observed with optogenetic FGFR1 multimerization depends on which terminus of FGFR1 the light-inducible oligomerization domain CRY2 is fused to, and whether this effect also occurs with other RTKs, we generated additional constructs: 1) Optodroplet-FGFR1, in which the Optodroplet module was fused to the N-terminus of FGFR1 **(Figure S1c,d)**; and 2) EphB2-optodroplet and optodroplet-EphB2, in which EphB2^46^ is another well-studied RTK **(Figure S1c,e)**. Quantification of ERK activity in cells expressing these constructs **(Figure S1f)** revealed similarly elevated p-ERK levels without light illumination, indicating that leaky activation is not dependent on the CRY2 fusion site (N- vs. C-terminal) and is not restricted to FGFR1. These findings suggest that basal ERK activation is a general property of CRY2-based optogenetic RTK constructs.

Together, these results indicate that CRY2-based RTK fusion proteins exhibit basal leaky ERK signaling, likely due to the intrinsic self-association propensity of CRY2. This effect is particularly pronounced in constructs containing the Optodroplet module. Membrane localization may further amplify clustering through two-dimensional confinement and increased local concentration^47^. Although blue-light stimulation enhances RTK clustering and ERK signaling output, the elevated basal activity observed in the absence of light may limit the utility of CRY2-based optogenetic systems in contexts requiring a tight input–output coupling. These findings highlight the need for alternative systems with improved control over RTK multimerization and more stringent regulation of downstream signal induction.

### Optimized rapamycin-inducible systems enable precise FGFR1 multimerization with minimal basal ERK activation

To circumvent the elevated basal activity observed with optogenetic FGFR1 constructs, we developed a novel chemically inducible system based on rapamycin-dependent heterodimerization of FKBP and FRB domains. Upon rapamycin addition, the FKBP–FRB pair rapidly associates with high affinity, enabling temporally precise induction of intracellular complex formation^48^. To promote higher-order clustering, we incorporated HOTag3 and HOTag6, two self-assembling peptide modules that form hexametric and tetrameric structures^26^. By fusing these elements to the intracellular domain of FGFR1, we designed a series of modular systems in which receptor multimerization could be tightly controlled, aiming to identify variants with low basal ERK activation in the absence of chemical stimulation **(Figure 2a)**.

**Figure 2.**
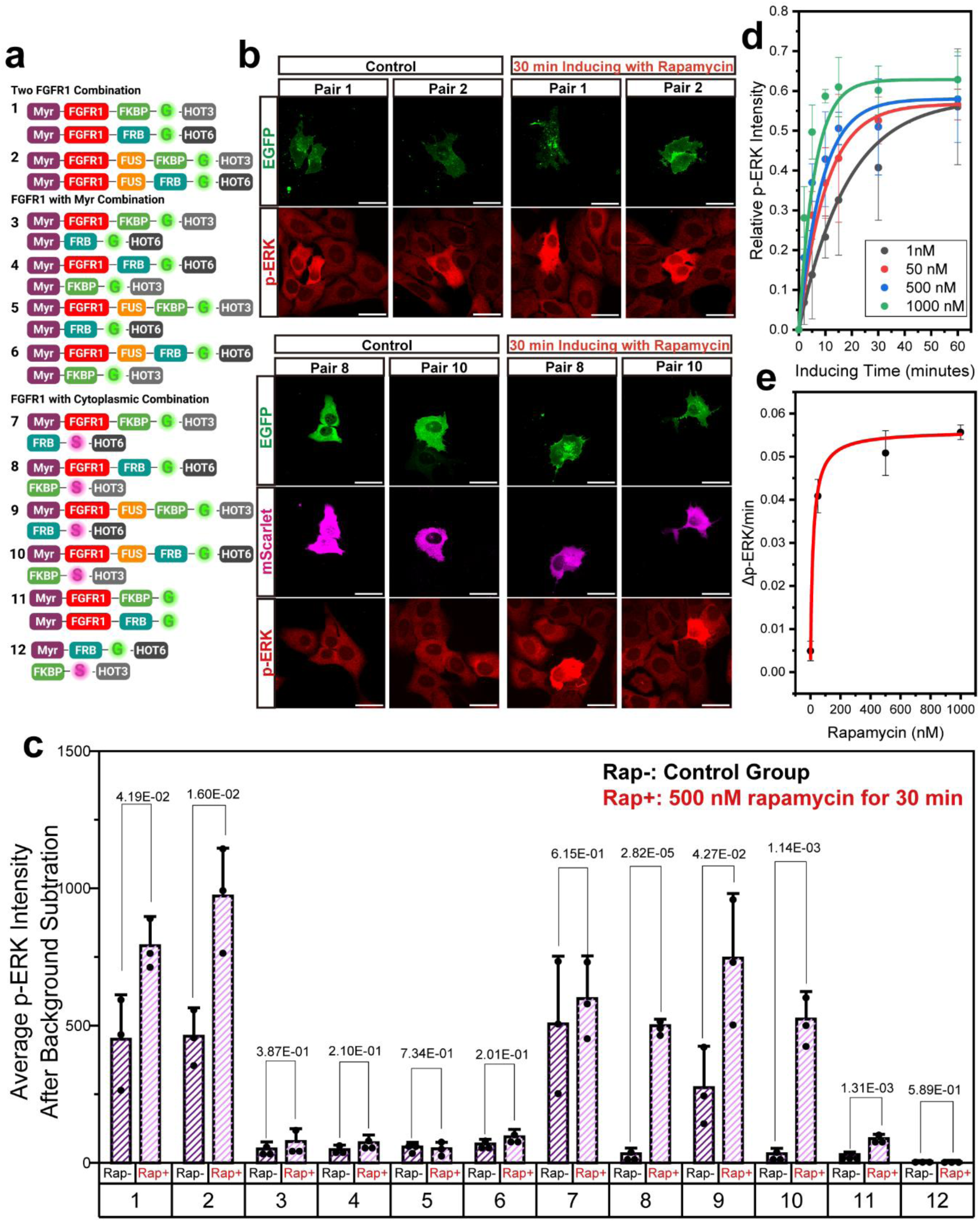
Systematic design and screening for rapamycin-inducible FGFR1 clustering construct with minimal basal ERK activation. **(a)** Schematic diagrams of all ten construct pairs (Pairs 1–12) **(b)** Representative fixed-cell confocal immunofluorescence images of U2OS cells expressing each construct pair. Scale bars, 10 μm. (**c)** Quantification of average total p-ERK fluorescence **(d)** Time-course of rapamycin-induced ERK activation for Pair 8. **(e)** Dose–response relationship of rapamycin-induced ERK activation rate for Pair 8. Data were fitted using the Michaelis–Menten kinetic model, yielding an apparent K_m_ of ∼17 nM.

In the initial construct designs (Pair 1 and Pair 2), both interacting components consisted of membrane-anchored CytoFGFR1 fused to either FKBP-HOTag3 or FRB-HOTag6 **(Figure S3)**. Pair 2 incorporated an additional intrinsically disordered FUS domain on both components to further enhance clustering propensity. Following rapamycin treatment for 30 min, duration that does not cause detectable cytotoxicity (**Figure S2a**), both pairs exhibited elevated levels of p-ERK, confirming activation of the ERK pathway **(Figure 2b, upper panels; Figure 2c)**. However, substantial basal ERK activity was also observed in the absence of rapamycin stimulation, indicating that membrane anchoring of both components, combined with multimeric scaffolds, was sufficient to promote chemical-independent receptor clustering.

To mitigate the basal activation, we designed a second set of constructs (Pairs 3–6) in which one of the two interacting components lacked the CytoFGFR1 domain but remained membrane-anchored via an N-terminal Myr motif **(Figure 2a; Figure S3, middle panels)**. This design aimed to reduce steric crowding while preserving rapamycin-inducible clustering capability. However, none of these pairs increased ERK phosphorylation upon rapamycin addition **(Figure S4a; Figure 3c)**, suggesting that they failed to form signaling-competent receptor assemblies. We speculate that tethering both components to the plasma membrane restricts conformational flexibility, thereby impeding the formation of productive higher-order oligomers^49^ and preventing the CytoFGFR1 domains from approaching closely enough to support signaling.

**Figure 3.**
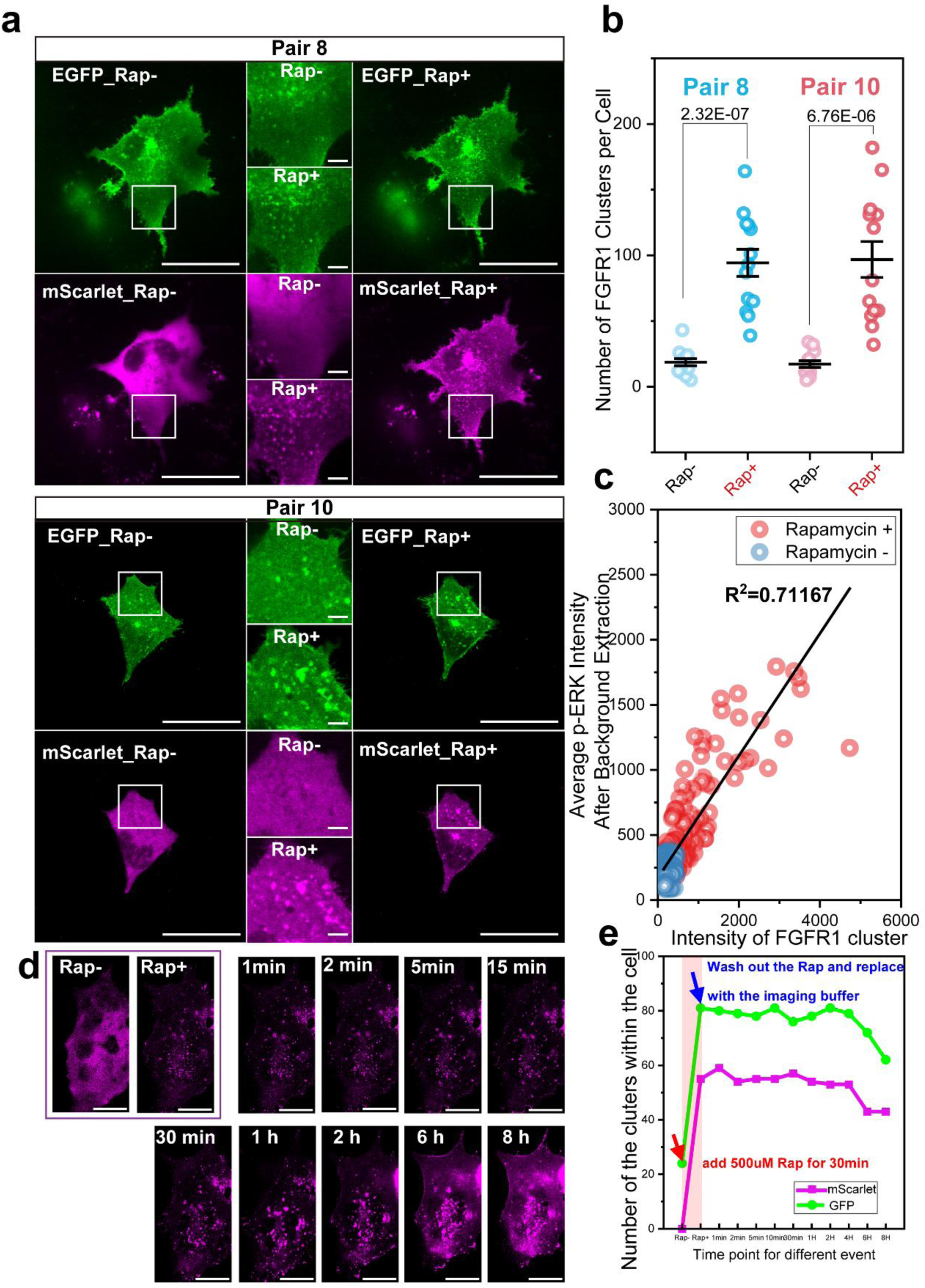
Rapamycin-induced FGFR1 clustering correlates with ERK pathway activation at single-cell level. **(a)** Representative live-cell confocal images of U2OS cells expressing construct pair 8 (top) or pair 10 (bottom). (**b)** Quantification of the number of clusters per cell (**c)** Correlation between total fluorescence intensity within the clusters and total p-ERK fluorescence intensity (R² = 0.712). **(d)** Time-lapse confocal imaging of U2OS cells expressing the optimized FGFR1 construct (Pair 8) under different stages of rapamycin stimulation. Cells were imaged in the absence of rapamycin (Rap–), immediately after addition of 500 nM rapamycin (Rap+), and at the indicated time points after rapamycin wash out. **(e)** Quantification of the cluster number over 8 hours.

To address the hypothesized factors that may have prevented rapamycin-induced clustering in Pairs 3-6, we implemented a third design strategy in which one interacting component was localized to the cytosol as a flexible bridging linker, while the other component remained a membrane-anchored CytoFGFR1 fusion (Pairs 7–10) **(Figure 2a; Figure S3)**. Among these constructs, Pairs 8 and 10 showed strong ERK activation upon rapamycin treatment with minimal basal signaling, producing robust increases in p-ERK levels **(Figure 2b, bottom panels; Figure 2c)**. In contrast, Pairs 7 and 9 displayed elevated p-ERK even without chemical stimulation **(Figure S4b; Figure 2c)**. Notably, in both pairs with high basal ERK background (i.e., Pairs 7 9), CytoFGFR1 was fused to HOTag3 (hexameric), a higher-valency domain with stronger self-association than HOTag6 (tetrameric). The increased basal ERK activity observed likely reflects enhanced spontaneous clustering driven by HOTag3. In the optimized pairs (8 and 10), HOTag3 is instead placed on the cytosolic bridging linker, where its higher valency enhances rapamycin-dependent higher-order assembly. These arrangements provides low basal signaling while enabling strong, stimulus-dependent higher-order assembly. To further confirm that HOTag-mediated higher-order assembly is required for both visible cluster formation and sufficient ERK activation, we generated Pair 11, where membrane-anchored CytoFGFR1 was fused to either FKBP or FRB without a HOTag domain (Figure **2a,c** and **Figure S5**). Pair 11 failed to form discernible clusters following rapamycin treatment, and the resulting p-ERK increase was minimal, falling below the threshold for robust activation. As an additional control, we removed the CytoFGFR1 domain from Pair 8 to generate Pair 12. Pair 12 displayed an increase in cluster numbers upon rapamycin treatment, yet p-ERK levels remained at baseline, demonstrating that ERK activation requires RTK signaling rather than oligomerization alone.

To assess the generalizability of our system, we replaced the CytoFGFR1 domain in Pair 8 with the intracellular domains of two additional RTKs, EphB2 and TrkB. Rapamycin-induced clustering of either receptor robustly activated ERK with minimal basal activity, mirroring the behavior of FGFR1 (**Figure S6**). These results indicate that our optimized platform can be readily extended to other RTKs.

Because blue-light illumination causes unwanted ERK activation in optogenetic systems (**Figure S2b**), we also evaluated whether rapamycin itself induces ERK signaling in our experimental window. Treatment with rapamycin (1-1000 nM) for durations ranging from 2 to 60 minutes did not significantly activate ERK (**Figure S2b**). These results confirm that our rapamycin-inducible RTK multimerization system further avoids the undesired inducer-dependent basal ERK activation observed in optogenetic approaches.

Next, to evaluate the tunability of the system, we used Pair 8 for kinetic characterization (**Figure 2d and 2e**). The apparent *K*_m_ for rapamycin is ∼17 nM, and the dynamic response range spans from low-pM level to ∼100 nM. These results indicate that the system is highly sensitive to rapamycin dosage. To ensure full activation in all subsequent experiments, we applied 500 nM rapamycin and induced cells for 30 min, a condition that consistently drives the system to its saturated equilibrium state.

Together, these results highlight the importance of tuning both spatial asymmetry and multimerization strength to achieve precise, chemically inducible RTK multimerization with minimal basal ERK activation, while allowing induced dosage to fine-tune the magnitude and kinetics of pathway activation.

### Inducible FGFR1 clustering drives proportional ERK activation

To directly visualize RTK clustering dynamics, we performed time-lapse live-cell imaging in cells expressing either Pair 8 or 10 **(Figure 3a)**. Upon rapamycin addition, both the EGFP-tagged FGFR1 component and the mScarlet-tagged cytosolic partner rapidly redistributed into discrete clusters that are colocalized in both color-channels. Quantitative analysis revealed a significant increase in cluster number after stimulation, confirming efficient RTK multimerization **(Figure 3b)**, demonstrating efficient and temporally precise induction of FGFR1 multimerization. To resolve these structures at higher spatial resolution, we performed 3D-STORM imaging on Pair 8 before and after rapamycin induction. STORM revealed a marked increase in FGFR1 cluster number **(Figure S7a-e)**, consistent with confocal maging(**Figure 3a,b**), and, single-molecule localization analysis showed substantially more localizations per cluster after induction, indicating higher molecular density **(Figure S7f)**. These observations confirm robust rapamycin-induced FGFR1 multimerization.

To investigate the relationship between receptor clustering and downstream ERK signaling strength, we examined how rapamycin-induced FGFR1 multimerization affected ERK activation at the single-cell level in cells expressing Pair 8 or 10. Immunofluorescence imaging on revealed substantial heterogeneity in both cluster density and p-ERK fluorescence intensity across individual cells. We quantified cluster intensity and p-ERK fluorescence intensity across a large population of transfected cells and observed that cells with higher levels of receptor clustering exhibited stronger ERK phosphorylation **(Figure 3c; Figure S8a, b)**. Linear regression analysis revealed a strong positive correlation between the fluorescence intensity of p-ERK with the EGFP signal from the cluster (R² =0.712), which the total EGFP signal (R²=0.382) and diffused EGFP signal (R²=0.349) do not show a significant linear relationship, indicating that ERK activation scales proportionally with the extent of receptor multimerization in our rapamycin-inducible system.

To assess the reversibility of rapamycin-induced RTK clustering, we performed extended time-lapse imaging for up to 8 hours following stimulation (**Figure 3d,e**). After rapamycin wash out, the pre-formed clusters persisted with cluster number unchanged over the 8-hour observation period, indicating that rapamycin-induced FGFR1 multimerization is effectively irreversible on the timescale of hours and generates highly stable receptor assemblies.

Together, these findings show that the rapamycin-inducible system enables rapid, spatially confined, and long-lasting FGFR1 multimerization, and that ERK signaling strength is determined by the extent of clustering rather than protein abundance. This supports a model in which receptor assembly state, rather than expression level, is the primary determinant of downstream signaling output.

### Rapamycin-inducible RTK system enables on-demand, background-free membrane skeleton disassembly

To determine whether ERK activation mediated by our system elicits functional downstream effects and to highlight the advantage of its background-free activation, we assessed the structural integrity of the spectrin-based submembranous cytoskeleton before and after stimulation. Sustained ERK activation has been shown to promote protease-dependent spectrin degradation and hence the disruption of spectrin-based membrane skeleton in various cell types, including neurons and epithelial cells^9^. We hypothesized that if rapamycin-induced FGFR1 clustering activates ERK in a temporally sustained manner, it should lead to disruption of spectrin-based membrane skeleton.

We focused on Pairs 8 and 10, which exhibited rapamycin-induced ERK activation with minimal basal ERK activity. Cells expressing each pair were treated with rapamycin for 30 minutes, fixed, and immunostained for endogenous βII-spectrin. For each imaging field of view (FOV), both transfected and neighboring untransfected cells were imaged to enable direct side-by-side comparison of cytoskeletal architecture within the same local microenvironment **(Figure 4a)**, thereby minimizing potential variability in cell density, substrate stiffness, and staining intensity across samples.

**Figure 4.**
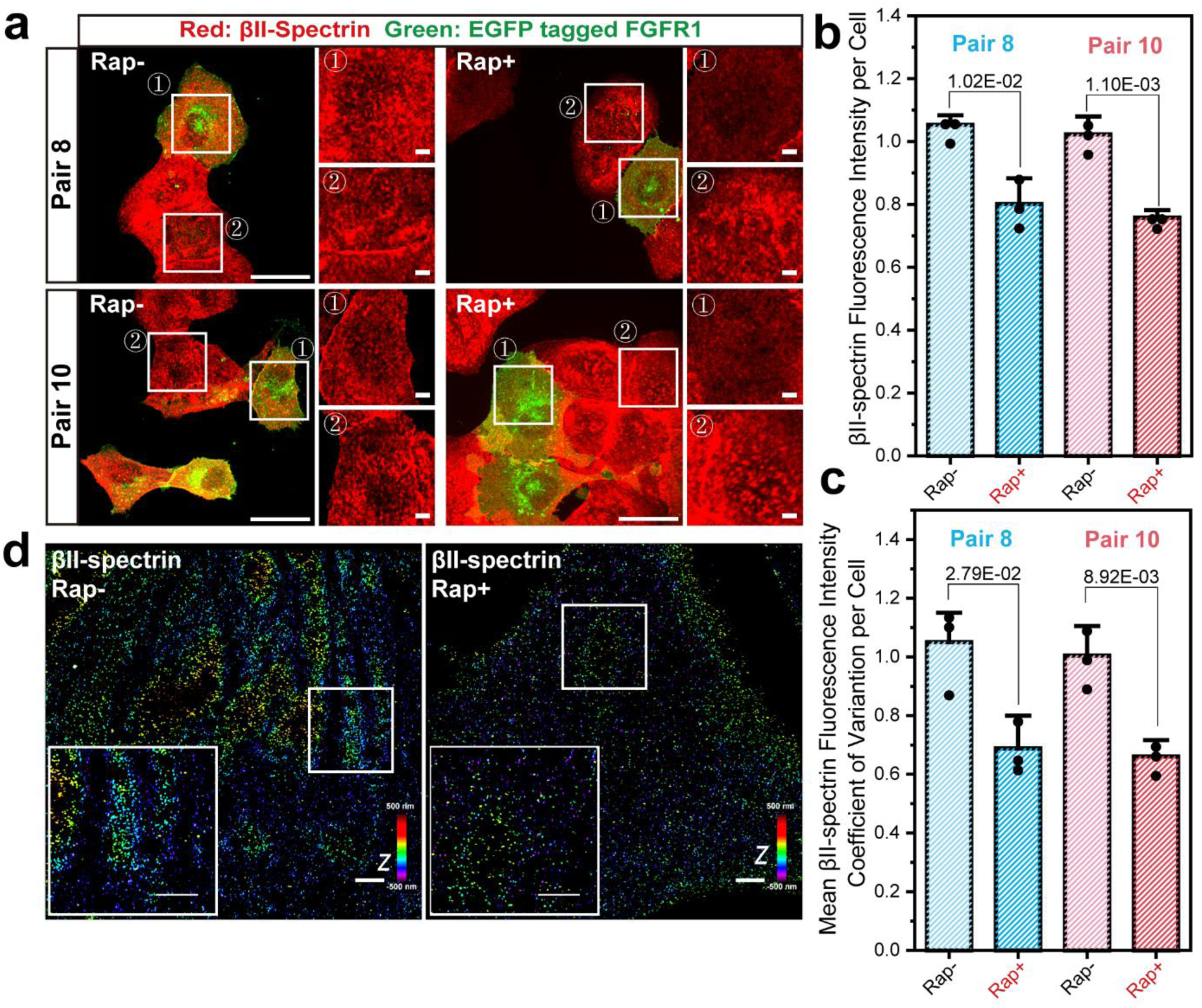
Controlled, background-free disassembly of the spectrin-based membrane skeleton via our rapamycin-inducible system. **(a)** Representative widefield immunofluorescence images of U2OS cells expressing Pair 8 (top) or Pair 10 (bottom). **(b)** Quantification of mean βII-spectrin fluorescence intensity per cell under each condition. **(c)** Mean coefficient of variation (CV) of βII-spectrin fluorescence intensity. **(d)** 3D-STORM super-resolution images of βII-spectrin in U2OS cells expressing Pair 8.

Without rapamycin, both transfected and untransfected cells expressing Pair 8 or 10 exhibited the expected spectrin immunostaining patterns, characterized by the puncta-like, heterogeneous islands of fluorescence located at the plasma membrane. This organization is consistent with patterns reported previously^50^. Following rapamycin stimulation, transfected cells displayed marked disruption of the membrane skeleton, where βII-spectrin fluorescence signal became diffuse, homogeneously distributed, and reduced in intensity, lacking the characteristic membrane enrichment and spatial heterogeneity. In contrast, neighboring untransfected cells retained intact spectrin architecture, indicating that the observed disassembly in transfected cells was specifically triggered by FGFR1 activation rather than rapamycin alone. By comparison, cells transfected with optogenetic constructs or other rapamycin-inducible FGFR1 pairs exhibited spectrin disruption even prior to stimulation **(Figure S9a)**, consistent with their elevated basal ERK activity.

To quantitatively assess the remodeling of spectrin membrane skeleton, we extracted two parameters from each cell: 1) mean βII-spectrin fluorescence intensity as a proxy for protein abundance **(Figure 4b)**; and 2) mean coefficient of variation (CV) of pixel intensity to reflect spatial patterning **(Figure 4c)**. Cells with intact spectrin architecture exhibited high CV values due to alternating regions of dense and sparse fluorescence, whereas disrupted networks displayed more homogeneous signal distribution and correspondingly lower CVs. In Pair 8, rapamycin-treated transfected cells exhibited a 25% reduction in spectrin intensity and a 30% decrease in CV, consistent with substantial cytoskeletal disassembly. Similar trends were observed in Pair 10. In both cases, untransfected cells from the same imaging fields showed unchanged intensity and CV values before and after rapamycin treatment **(Figure S9b)**, confirming that spectrin degradation was specifically linked to FGFR1 clustering and downstream ERK activation.

To further validate these observations at the nanoscale, we performed 3D-STORM imaging of βII-spectrin in cells expressing Pair 8 **(Figure 4d)**. Without rapamycin, spectrin formed elongated, isolated islands along the inner leaflet of the plasma membrane, and within each island, a lattice-like arrangement of βII-spectrin molecular clusters was observed, similar to spectrin organization previously reported in neuronal soma^51^ or red blood cells^52^. These elongated spectrin islands are consistent with exclusion patterns previously observed between the spectrin membrane skeleton and actin stress fibers in mouse embryonic fibroblasts (MEFs)^53,54^. After rapamycin treatment, these elongated spectrin islands disappeared, replaced by amorphous, isotropic, and markedly reduced density of βII-spectrin molecular clusters, consistent with cytoskeletal disassembly and loss of organized spectrin architecture.

Together, these results demonstrate that rapamycin-induced FGFR1 clustering leads to functionally relevant ERK activation, driving spectrin degradation and cytoskeletal remodeling. Notably, our chemical-inducible RTK platform shows no basal undesired induction compared to the existing inducible RTK systems. The integration of downstream functional readouts highlights the utility of this platform for probing how graded signal input strength governs cytoskeletal architecture.

### Rapamycin-inducible RTK system enables on-demand, background-free activation of transcription factors

Having demonstrated that our system can control disassembly of the spectrin-based membrane skeleton, we next explored its broader applicability in modulating other ERK-regulated cellular processes. As ERK phosphorylates and regulates the nuclear translocation of numerous transcription factors^55^, we focused on two well-characterized ERK-responsive effectors: 1)STAT3, which mediates proliferation, differentiation, and cellular stress response downstream of diverse RTKs^56^; and 2)CREB, which governs survival, plasticity, and cytoskeletal organization^57^..

Rapamycin treatment caused a robust increase in p-STAT3 and p-CREB signals in cells expressing Pair 8 or 10 **(Figure 5)**, while unstimulated cells maintained low basal levels comparable to adjacent untransfected cells, reflecting the minimal background activity of our system. In contrast, cells expressing optogenetic constructs or Pair 1 showed substantial pre-activation even before stimulation **(Figure S10)**. These results demonstrate that chemically induced FGFR1 clustering is sufficient to drive not only proximal ERK phosphorylation but also downstream nuclear activation of key transcription factors. Importantly, this occurs without basal ERK activity, confirming that our system effectively recapitulates both cytoplasmic and nuclear RTK signaling in a tightly controlled, inducible manner.

**Figure 5.**
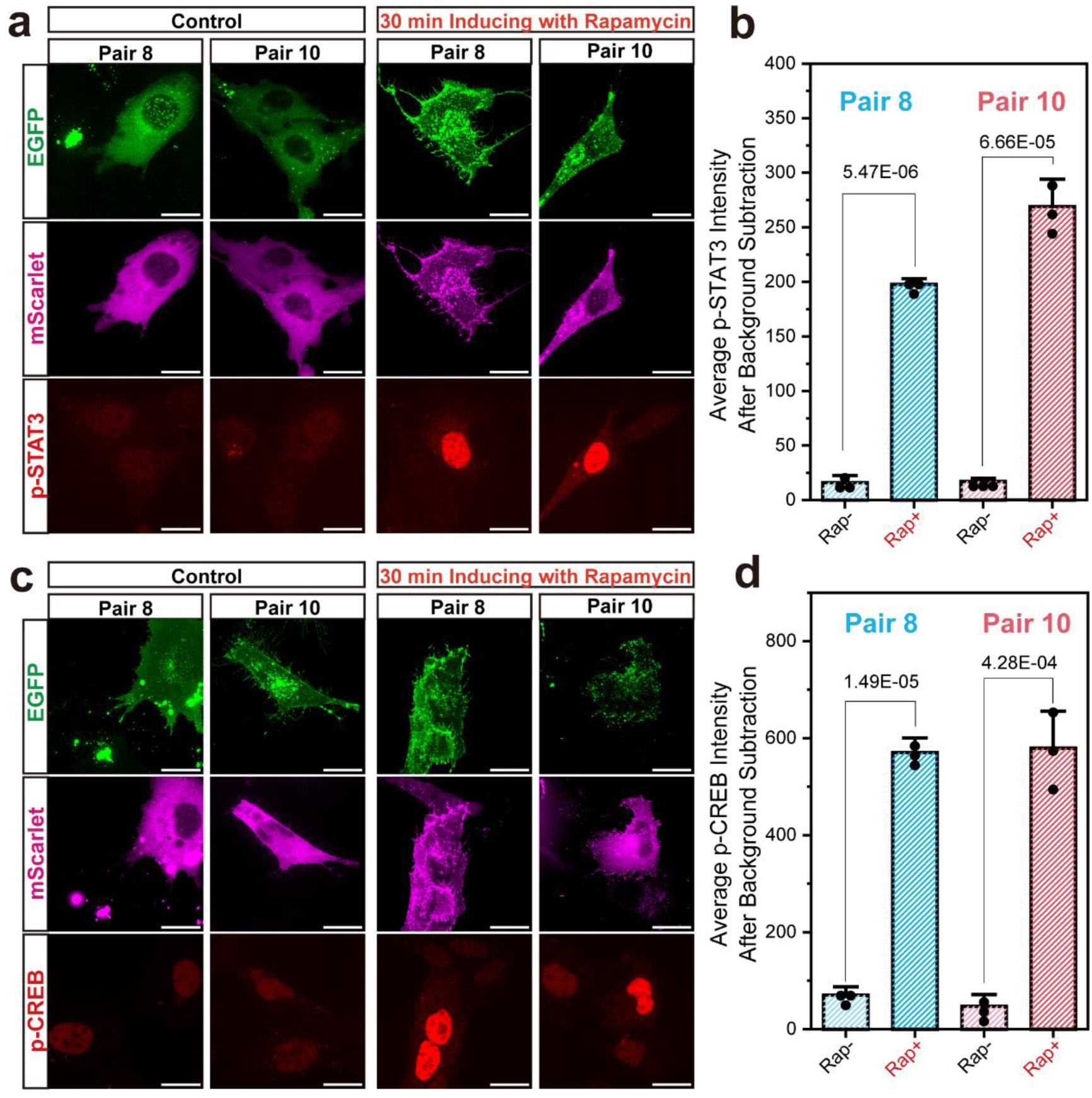
Chemically induced FGFR1 clustering activates ERK-dependent nuclear transcriptional programs. (a, b) Quantification of p-STAT3 fluorescence intensity in cells expressing FGFR1 clustering constructs (Pair 8 and Pair 10) before and after rapamycin treatment (250 nM, 15 min). Representative confocal images (a) and corresponding quantification (b). (c, d) Same as (a,b) but for p-CREB instead of STAT3.

### Working model linking RTK multimerization to downstream ERK-regulated cellular processes and spectrin degradation

To summarize our findings and highlight the modularity and biological relevance of synthetic RTK clustering, we developed a mechanistic model comparing optogenetic and chemically inducible systems **(Figure 6).**

**Figure 6.**
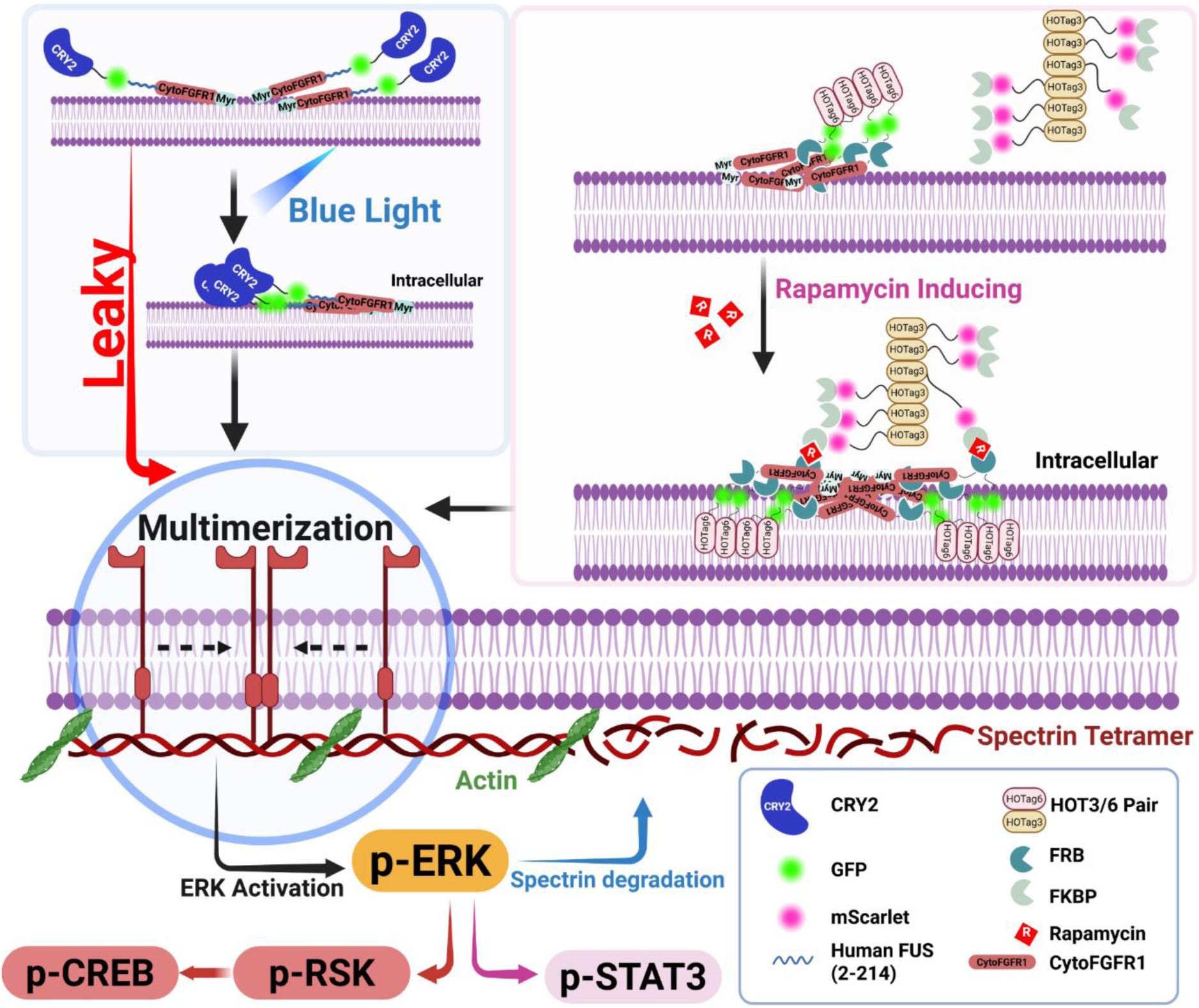
Mechanistic model comparing optogenetic and chemically inducible RTK clustering systems. Created in BioRender. Zheng, Y. (2025) https://BioRender.com/gxkzrjo

On the left, CRY2-based optogenetic FGFR1 constructs oligomerize upon blue-light stimulation to induce receptor clustering and ERK activation. However, intrinsic self-association of CRY2, enhanced in high-expression contexts or by oligomerization-promoting mutations (CRY2olig) and intrinsically disordered domains (e.g., FUS), leads to substantial basal clustering and ERK activation even without light. This leakiness reduces the dynamic range and limits precise input–output control.

On the right, the rapamycin-inducible system leverages FKBP–FRB heterodimerization to bring multimerization domains (HOTag3 or HOTag6) into proximity, driving FGFR1 clustering with minimal basal ERK activity. Spatial asymmetry (e.g., pairing membrane-localized FGFR1-FRB-HOTag6 with cytosolic HOTag3-FKBP) enables robust, tunable ERK activation and precise temporal control, with remarkable minimal basal ERK activation.

The model integrates downstream ERK-dependent effects across cellular compartments. In the cytoplasm, ERK activity drives βII-spectrin degradation and cortical cytoskeletal remodeling, as observed by widefield and 3D-STORM imaging. In the nucleus, ERK phosphorylates transcription factors STAT3 and CREB, regulating gene programs associated with proliferation, differentiation, survival, and structural remodeling. Importantly, our platform enables precise, tunable control of these compartment-specific processes while maintaining minimal basal activation. Overall, this model highlights the design flexibility and high signaling fidelity of chemically inducible clustering platforms, providing a versatile framework for dissecting how spatial and valency parameters encode biological responses.

## CONCLUSIONS

Precise modulation of RTK clustering remains a central objective in cell signaling research, as receptor multimerization serves as a key regulatory node linking extracellular ligand engagement to downstream biochemical and mechanical outputs^4^. In this study, we developed and characterized a chemically inducible system for RTK clustering based on rapamycin-induced FKBP-FRB heterodimerization^58,59^, in combination with the modular multimerization domain HOTag3 and HOTag6^25^. Systematic construct design and functional analyses demonstrate that this platform enables tightly controlled, low-background activation of the ERK signaling cascade. ERK activation through this inducible clustering system produces functional downstream effects ranging from nanoscale cortical cytoskeletal remodeling, including βII-spectrin disassembly, to nuclear activation of transcription factors STAT3 and CREB. These findings confirm that our synthetic clustering system faithfully recapitulates both cytoplasmic and nuclear RTK outputs while maintaining minimal basal activity, allowing precise and physiologically relevant control over signaling dynamics.

Comparison with CRY2-based optogenetic systems^60^ highlights key advantages: whereas CRY2 oligomerization often induces substantial basal ERK activity, our rapamycin-inducible constructs achieve minimal background signaling while retaining robust, tunable activation, providing superior dynamic range and signaling fidelity. While the system relies on the essentially irreversible FKBP–FRB interaction, future iterations could incorporate reversible dimerization modules to expand temporal control. The platform is also readily generalizable to other RTKs and membrane-associated signaling complexes, enabling the study of how clustering geometry, valency, and localization influence signaling specificity and temporal dynamics. Coupling this system with spatially restricted chemical inducers could allow subcellular control comparable to optogenetic tools, while preserving low basal activity.

Beyond single-cell applications, this system is compatible with multicellular and organoid models, where controlled RTK clustering could probe morphogen gradients, cell–cell communication, and tissue-level patterning. The observation that synthetic FGFR1 clustering drives spectrin disassembly also suggests utility for studying signal-to-mechanical feedback loops with implications for epithelial integrity, neuronal polarity, and cancer cell plasticity. In summary, this study establishes a chemically tunable and robust platform for RTK clustering that enables precise, on-demand control of receptor organization with minimal basal activation. This system allows dissection of the causal relationships between clustering, signaling strength, and downstream cellular responses. Its modular design and high signaling fidelity make it a versatile tool for probing and engineering cellular signaling networks with molecular precision.

## ASSOCIATED CONTENT

The following Supporting Information is available free of charge at ACS Website: Additional experimental data and analyses, including optimization of optogenetic and chemical-inducible FGFR1 clustering designs; ERK activation quantification; construct schematics; rapamycin cytotoxicity assays; STORM super-resolution reconstructions; single-cell signaling correlations; spectrin degradation imaging; STAT3 and CREB downstream activation; and expanded methods and references.

## AUTHOR INFORMATION

## Author Contributions

R.Z. conceived the project. Y.Z., J.F., and R.Z. designed experiments, interpreted the data and wrote the manuscript. Y.Z., J.F., A.C. performed experiments. Y.Z. and J.F. analyzed the data. R.Z. acquired funding and supervised the project.

## ACKNOWLEDGMENT

This work was supported by National Institute of General Medical Sciences (R35GM142973), the Life Sciences Research Foundation, and the startup fund from the Pennsylvania State University provided to R.Z.

## For Table of Contents Only

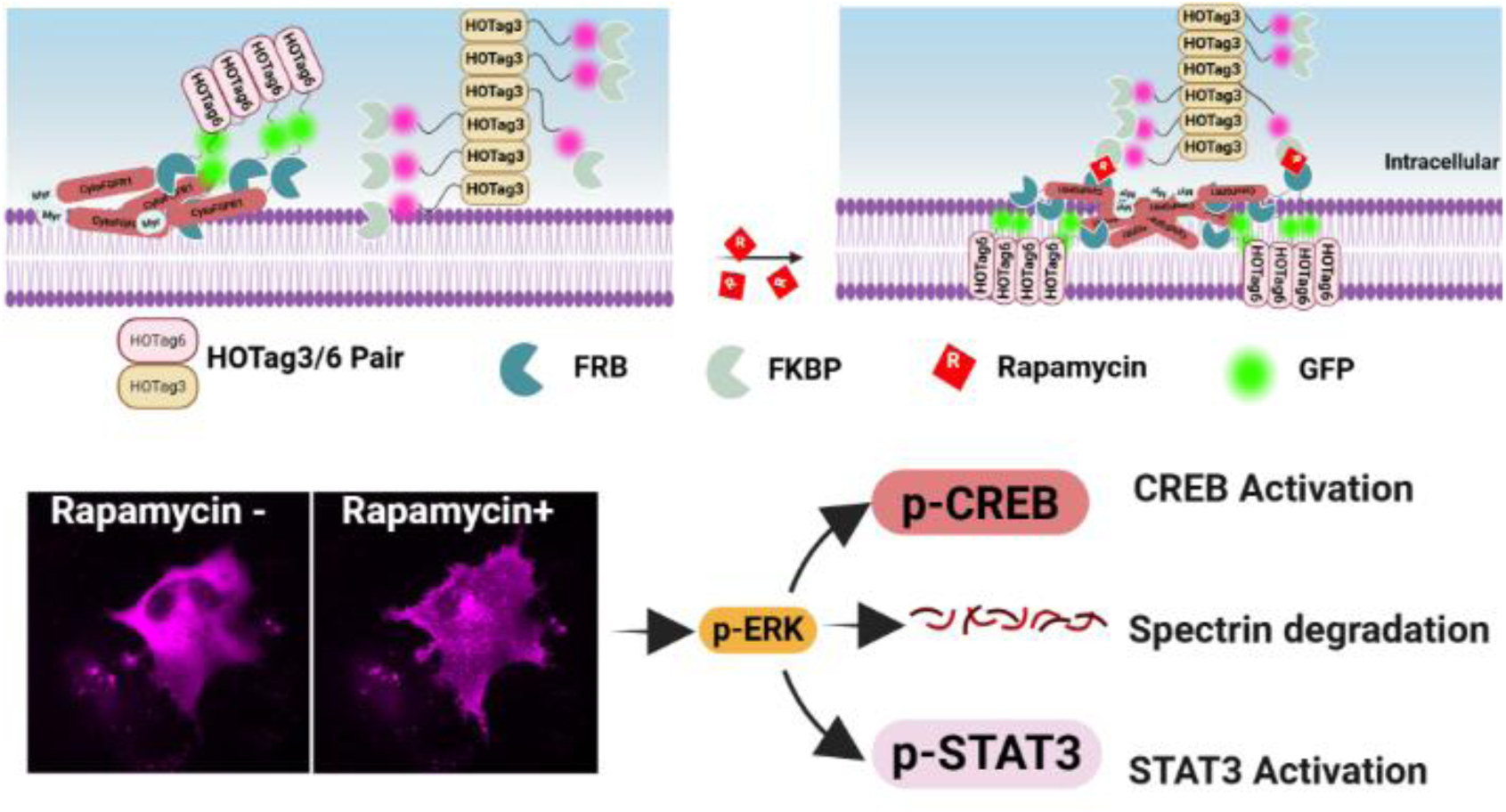

## Supporting Information

**Figure S1.**
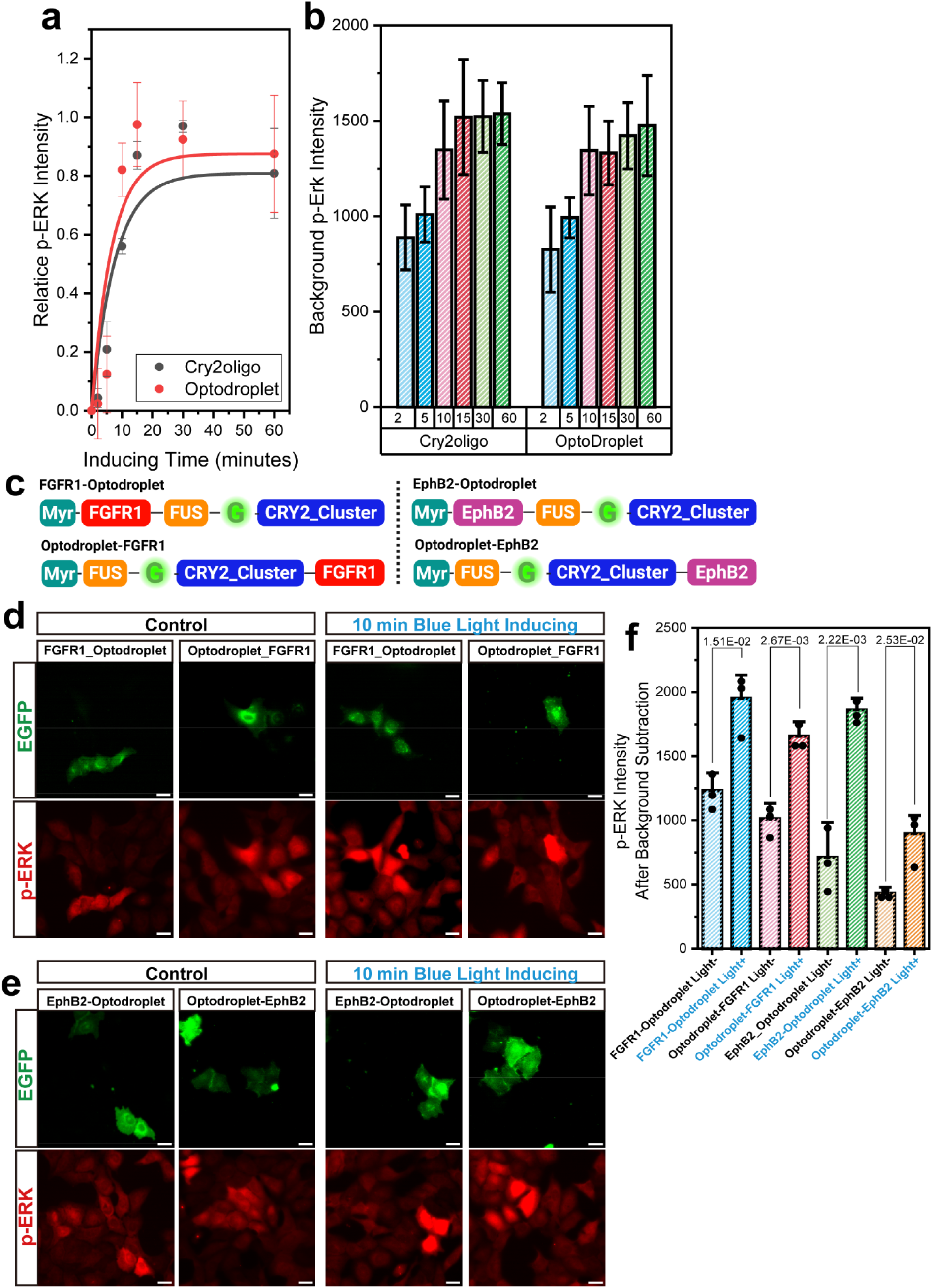
Leaky ERK activation is independent of the optoDroplet module’s N- or C-terminal position on RTK **(a)** Kinetic curves of relative p-ERK activation following blue-light stimulation for CRY2oligo and Optodroplet. Both constructs show rapid activation within the first 10–15 min, after which p-ERK levels plateau, defining the saturation window for optogenetic induction. **(b)** Quantification of basal p-ERK intensity in untransfected U2OS cells at the same sample with cell expressing FGFR1-CRY2oligomer (CRY2oligo) or FGFR1-Optodroplet constructs under different blue-light exposure durations (2, 5, 10, 15, 30, and 60 min). Bars represent mean ± s.d. from *n* = 3 biological replicates. **(c)** Schematic diagrams of FGFR1 and EphB2 constructs with the optoDroplet module attached at the N- or C-terminus of RTK. **(d) and (e)** Representative confocal immunofluorescence images of U2OS cells expressing FGFR1/EphB2-Optodroplet and Optodroplet-FGFR1/EphB2 constructs. All constructs were tagged with EGFP (green) and assessed for ERK pathway activation using immunostaining for p-ERK (red) under dark (control) and light-inducing conditions. Scale bars, 20□μm**. (f)** Quantification of average total p-ERK fluorescence intensity in transfected cells under control and light-stimulating conditions. Fluorescence intensities were background-subtracted using signals from neighboring untransfected cells and presented as mean ± s.d. from n□=□3 biological replicates. Individual data points represent averaged values from independent imaging fields.

**Figure S2.**
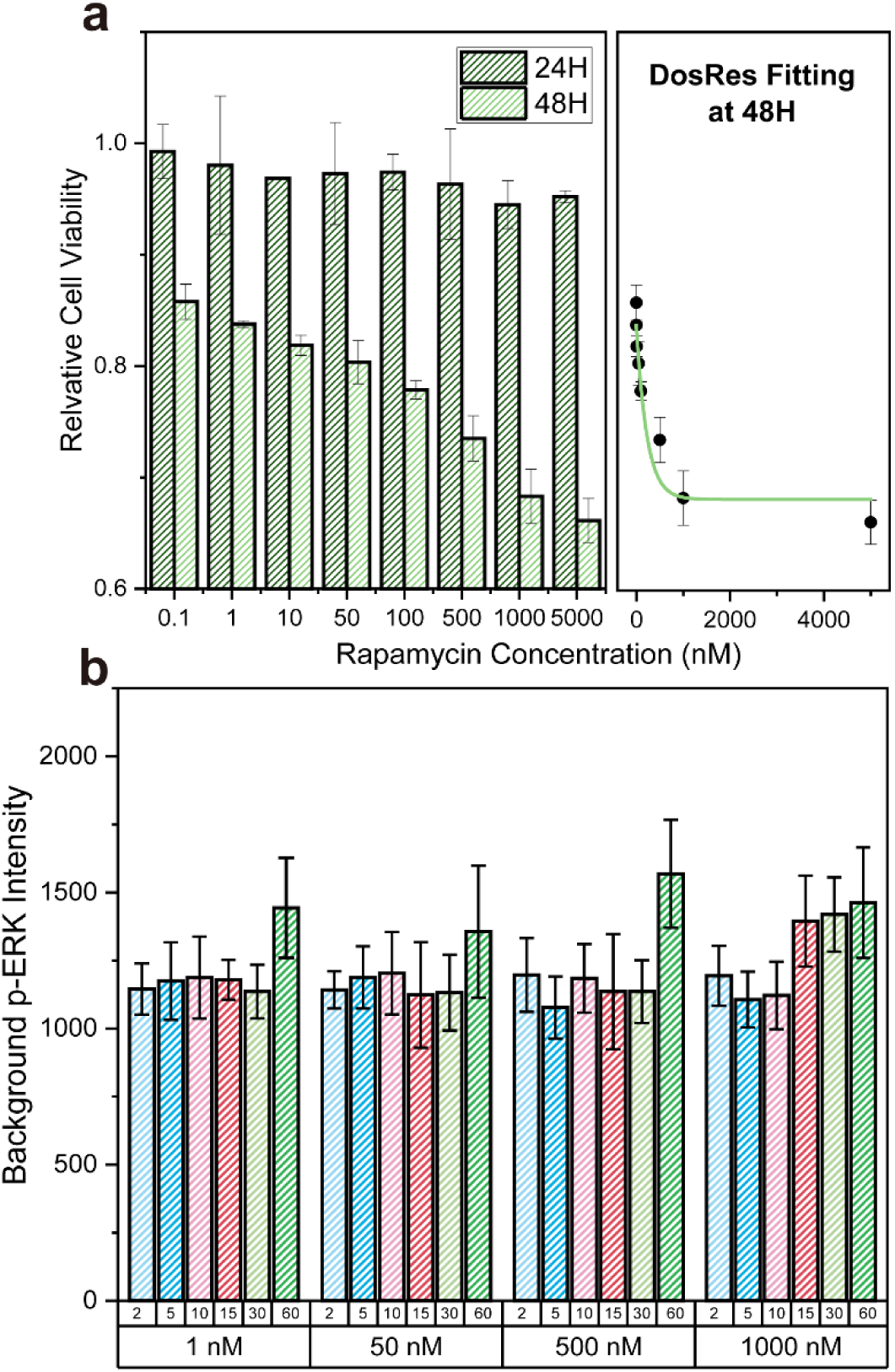
Rapamycin cytotoxicity and basal ERK activity across a range of rapamycin concentrations. **(a)** Cell viability measurements of U2OS cells treated with increasing concentrations of rapamycin (0.1–5000 nM) for 24 h and 48 h, using the MTT assay. Viability was normalized to untreated controls. A dose–response curve (right) was fitted to the 48 h data, showing minimal cytotoxicity below ∼1000 nM. These results indicate that short-term rapamycin stimulation, on the order of minutes to several tens of minutes as used in our clustering experiments, is unlikely to compromise cell viability. **(b)** Quantification of basal p-ERK levels in U2OS cells expressing the indicated constructs (color-coded) following 1, 15, 30, and 60 min exposure to rapamycin at 1, 50, 500, or 1000 nM. p-ERK intensities were background-subtracted using signals from untransfected neighboring cells. Bars represent mean ± s.d. Across all doses, rapamycin alone did not induce detectable ERK activation in the absence of FGFR1 clustering, confirming low off-target activity.

**Figure S3.**
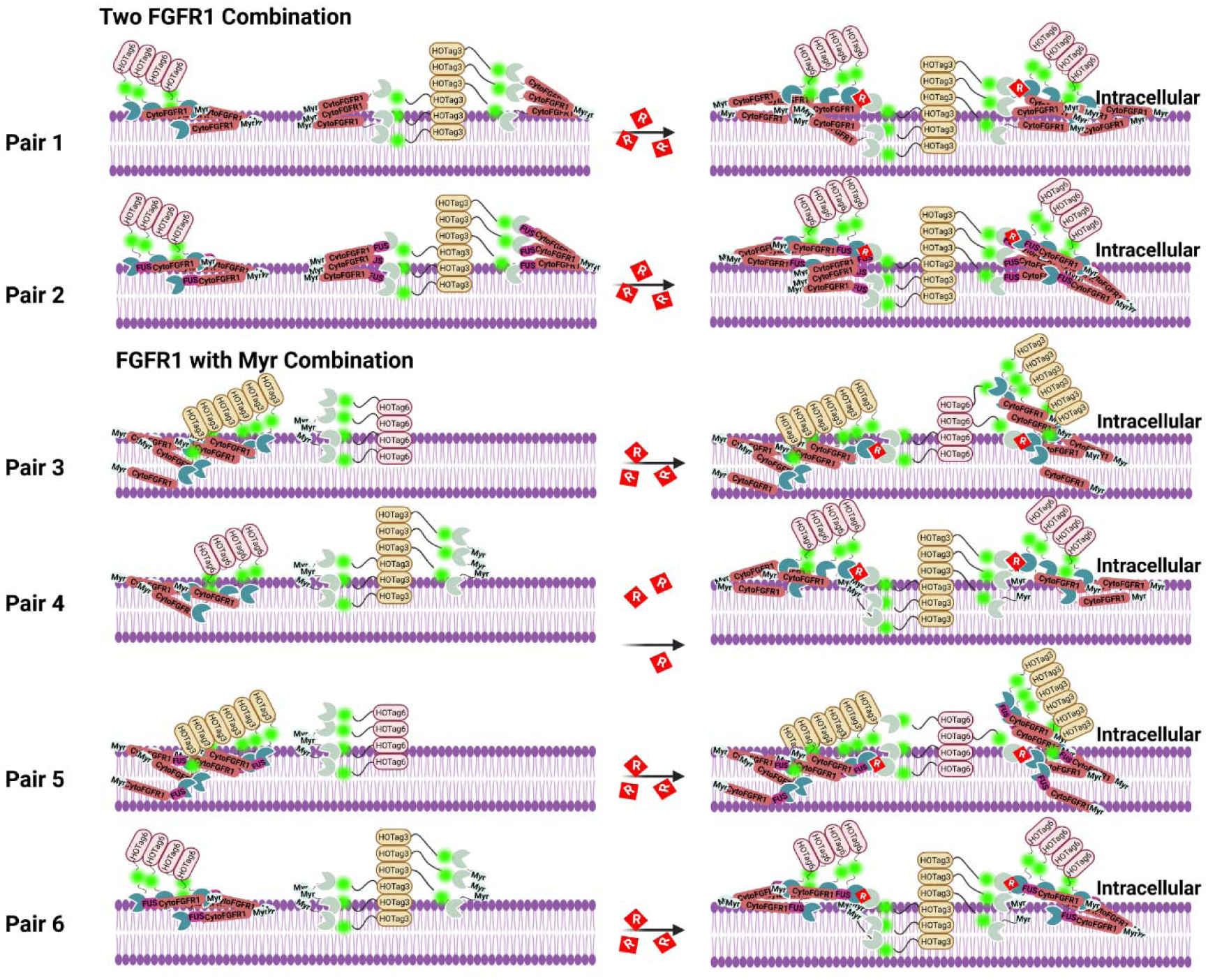
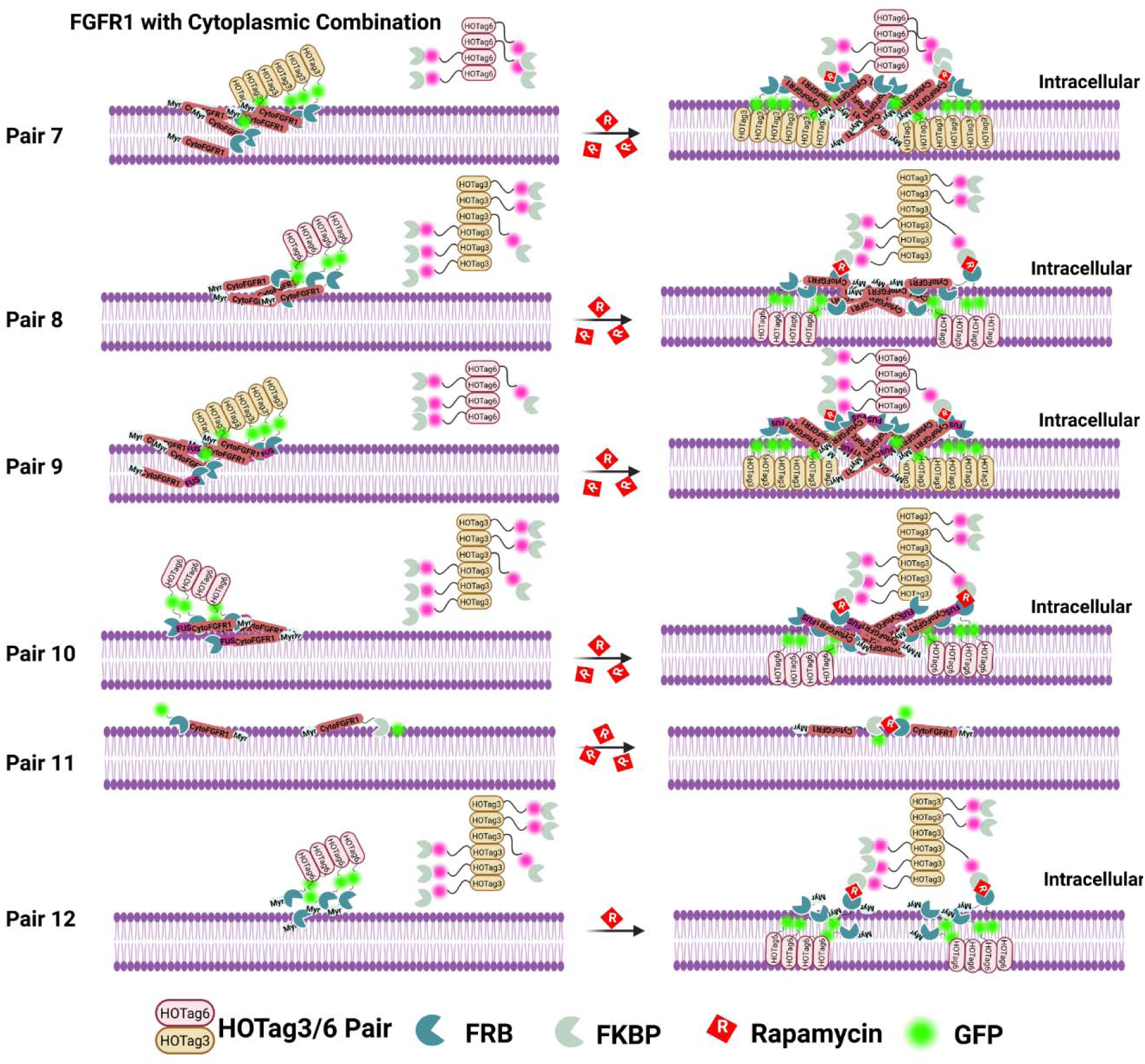
Schematic diagrams of three chemical-inducible RTK activation strategies. Mechanistic illustrations of three classes of chemically inducible FGFR1 clustering designs. All systems utilize rapamycin-induced FKBP–FRB heterodimerization to drive higher-order multimerization coupled to HOTag3 and HOTag6 coiled-coil domains. Membrane-tethered designs (Pairs 1–2), in which both FGFR1 fusion components are anchored to the plasma membrane and contain multimerization domains. These configurations promote strong receptor clustering but exhibit relatively high basal activity, potentially resulting from the high effective concentration of FGFR1 at the two-dimensional membrane. Asymmetric membrane–membrane designs (Pairs 3–6), in which one fusion construct lacks the FGFR1 kinase domain but remains membrane-anchored. These constructs reduce basal clustering but show weak ERK activation upon stimulation, potentially due to constrained clustering geometry preventing the higher-order FGFR1 multimerization. Asymmetric membrane–cytosolic designs (Pairs 7–10), where FGFR1 fusion constructs are membrane-localized while interacting partners reside in the cytosol. Rapamycin induces interfacial assembly at the membrane, enabling tunable clustering with minimal background activity. Optimal signaling output is achieved with balanced multimerization strength and spatial separation. Created in BioRender. Zheng, Y. (2025) https://BioRender.com/5i7pvbv

**Figure S4.**
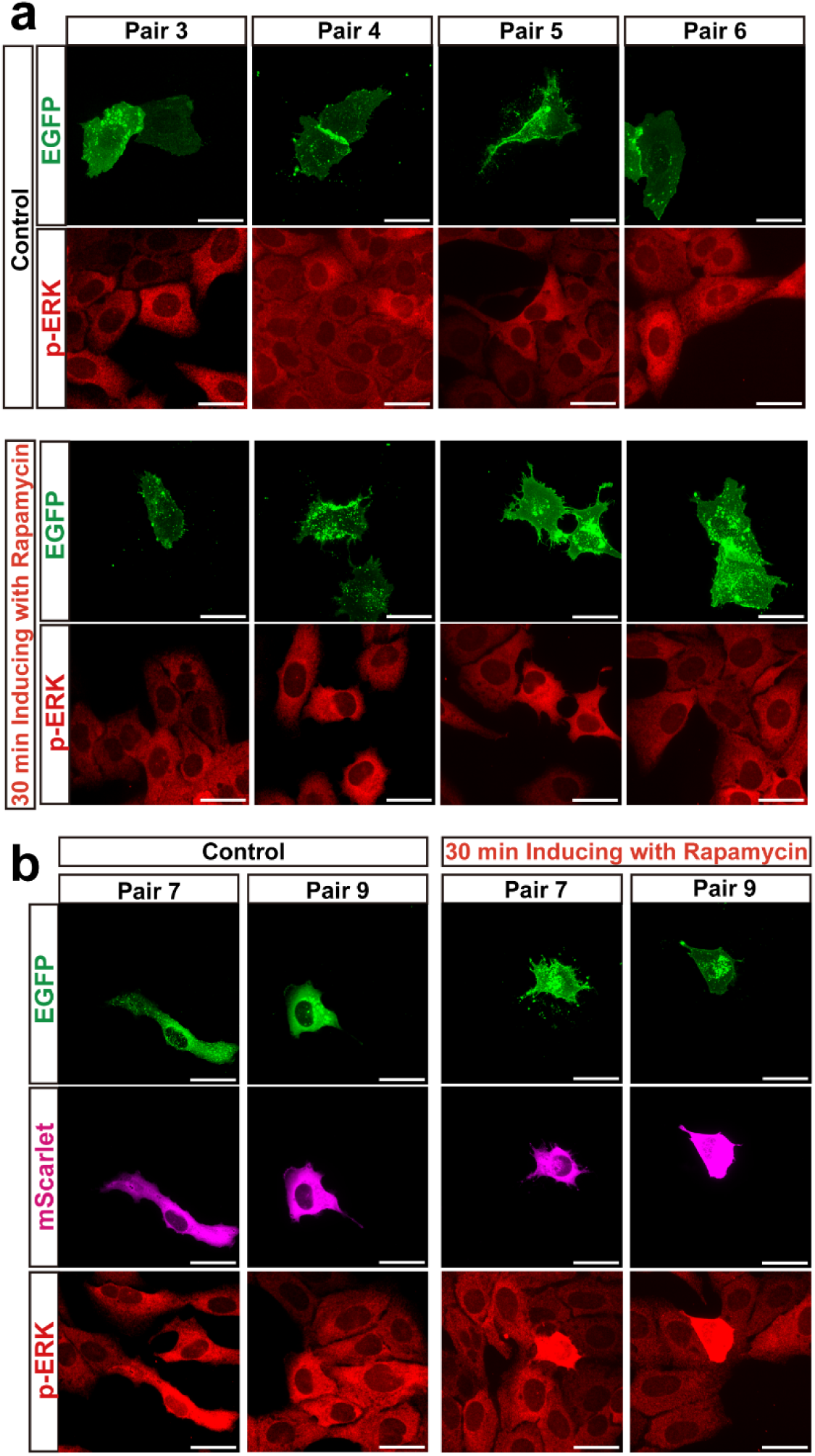
ERK activation of additional rapamycin-inducible designs. U2OS cells were transfected with the indicated FGFR1 constructs from screening Pairs 3-6, 7 and 9, and imaged under basal conditions or following 30-minute stimulation with 500□nM rapamycin. Each construct was tagged with EGFP (green) to visualize expression and clustering behavior. ERK activation was assessed via immunostaining for p-ERK (red). Cells expressing Pair 7 and Pair 9 also included mScarlet-tagged components (magenta) to distinguish membrane-localized and cytosolic modules. Constructs showed variable levels of basal ERK signal and inducible ERK activation depending on clustering configuration and multimerization strength. Scale bars, 10□μm

**Figure S5.**
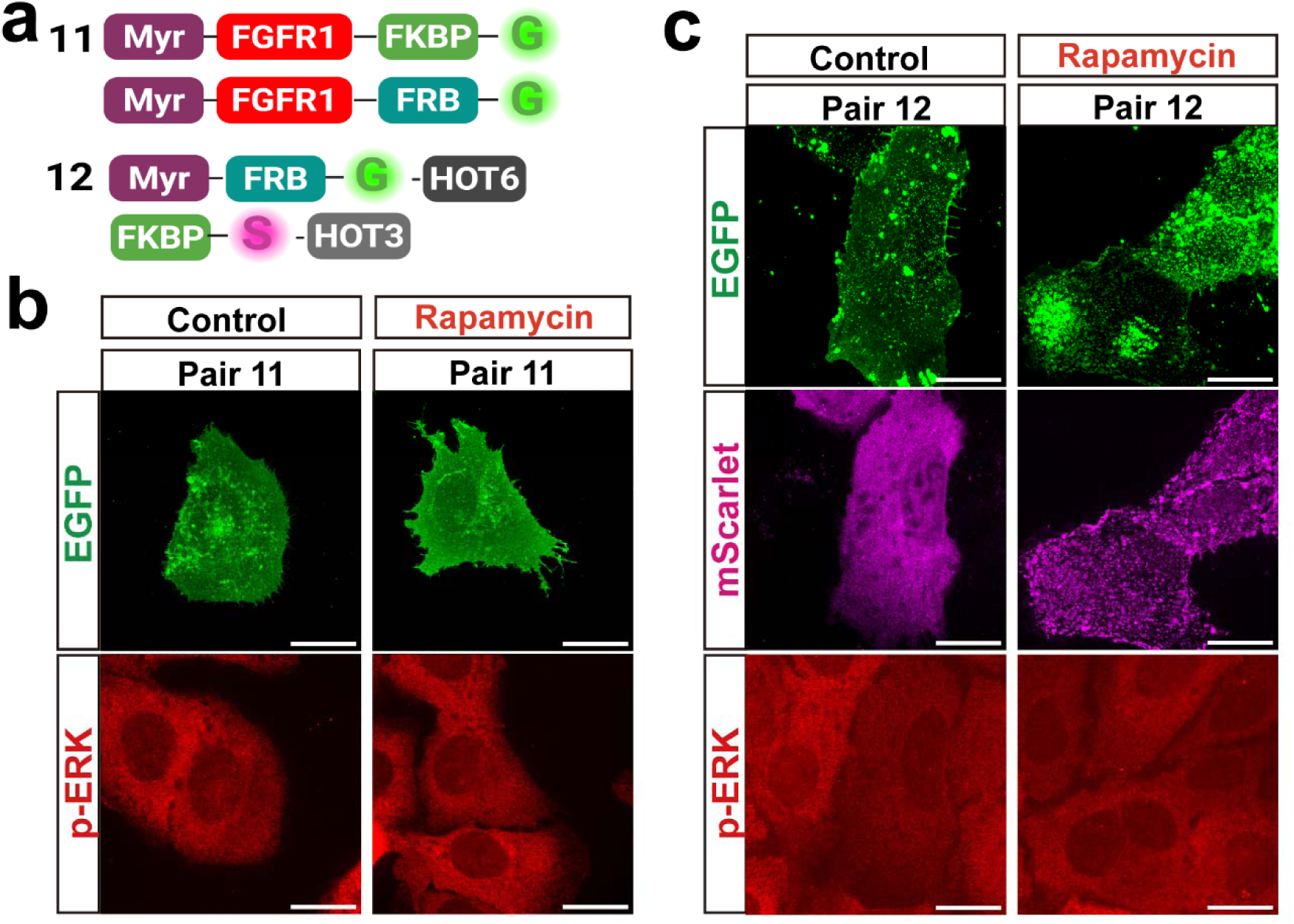
Evaluation of backbone constructs (Pairs 11 and 12) lacking FGFR1 or HOTag domains for clustering and ERK activation. **(a)** Schematic diagrams of Pair 11 and Pair 12 constructs used to test whether the control constructs without HOTags or without FGFR1 are capable of forming detectable clusters or activating downstream ERK signaling. **(b)** Representative confocal images of U2OS cells expressing Pair 11, under control and rapamycin-treated conditions (500 nM, 30 min). EGFP and mScarlet reporters indicate construct expression and subcellular localization. No discernible clustering or p-ERK elevation was observed upon rapamycin treatment, indicating minimal background activity of the Pair 11 backbone. **(c)** Representative images of U2OS cells expressing Pair 12 under the same conditions. Although rapamycin induced visible clustering due to the higher-order assembly modules HOTag3 and HOTag6, p-ERK levels remained comparable to untreated controls. These results demonstrate that the backbone constructs alone—even those capable of multimer formation—do not trigger ERK activation, confirming that receptor fusion is required for productive signaling.

**Figure S6.**
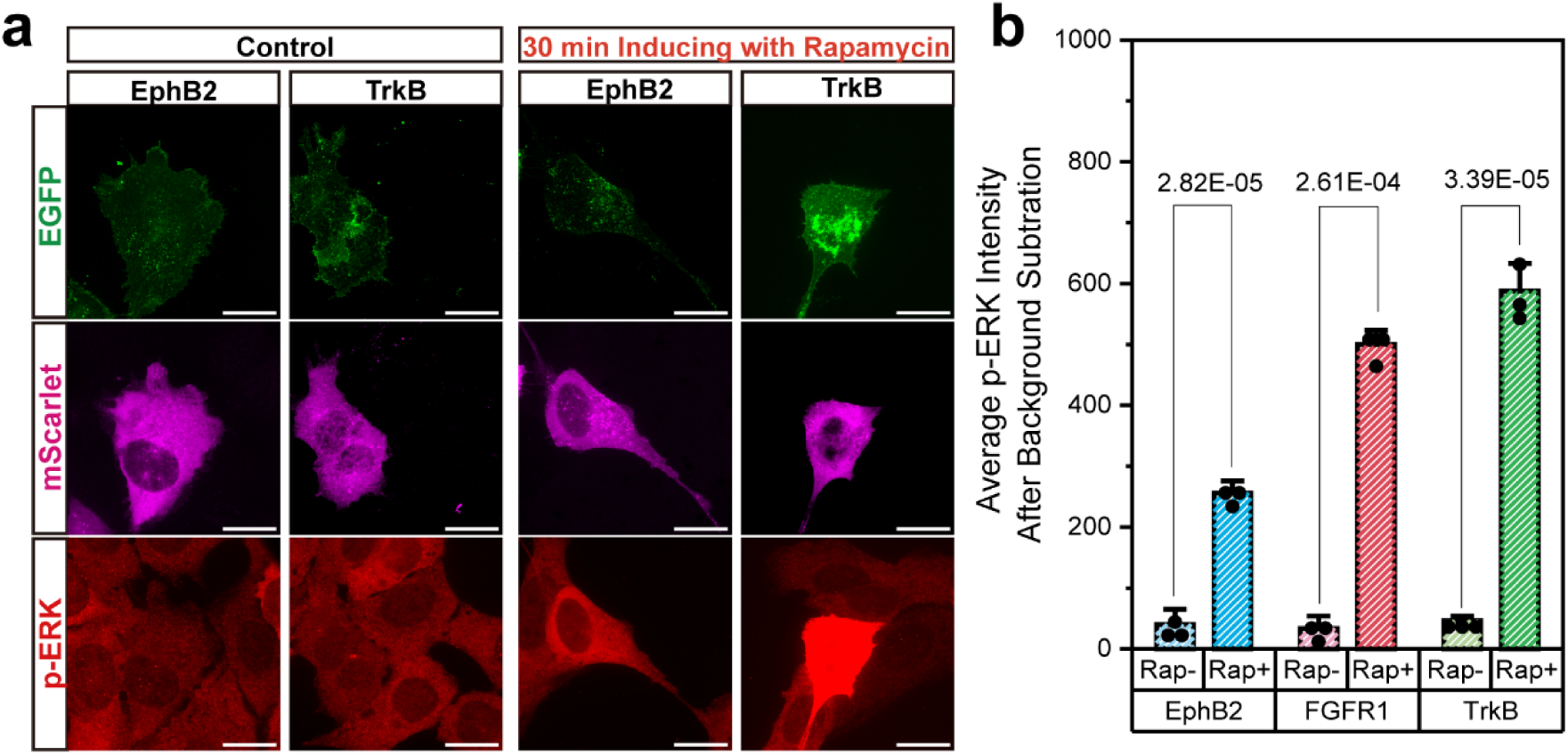
Generalizability of the chemical-inducible clustering system to alternative RTKs (EphB2 and TrkB). **(a)** Representative confocal images of U2OS cells expressing Pair 8 constructs in which the FGFR1 intracellular domain was replaced with the intracellular domain of either EphB2 or TrkB. Cells were imaged under control conditions or after rapamycin treatment (500 nM, 30 min). EGFP and mScarlet channels indicate expression and membrane localization of each RTK fusion pair. In both EphB2- and TrkB-based systems, rapamycin induced robust receptor clustering comparable to FGFR1. **(b)** Quantification of p-ERK levels across the three RTK types (EphB2, FGFR1, and TrkB) with or without rapamycin treatment. For each receptor, rapamycin stimulation resulted in a significant increase in p-ERK intensity after background subtraction, demonstrating that the chemically inducible Pair 8 design reliably activates ERK downstream of diverse RTKs. These results highlight the modularity and broad applicability of our clustering platform for reprogramming signaling across multiple receptor systems.

**Figure S7.**
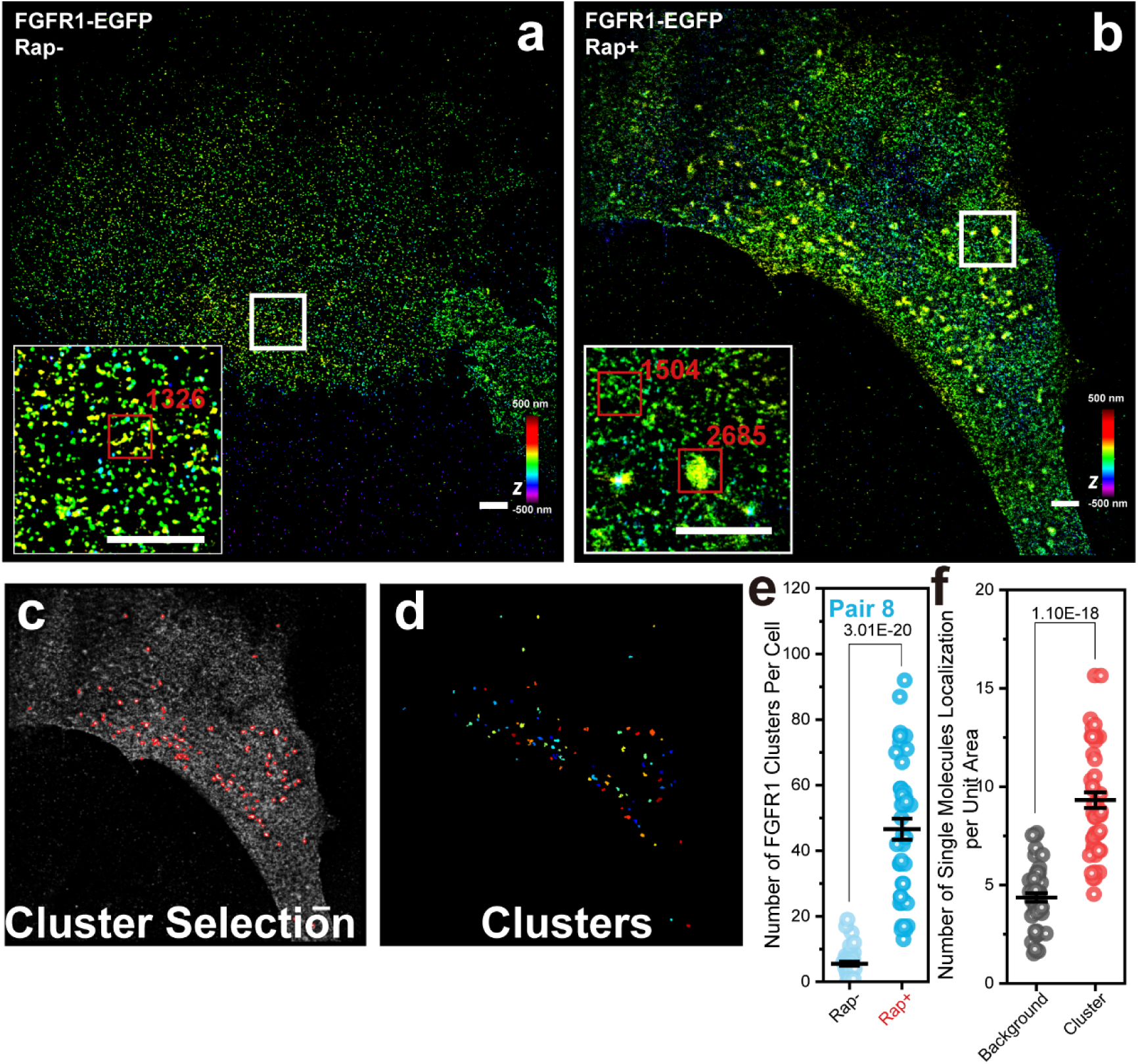
3D-STORM analysis of nanoscale FGFR1 clustering before and after rapamycin induction. **(a, b)** 3D-STORM reconstructions of FGFR1-EGFP labeled with AF647-conjugated anti-GFP antibody in U2OS cells expressing Pair 8, imaged under control conditions (Rap–) and after rapamycin treatment (Rap+, 500 nM, 30 min). The *z*-position of each localization is color-coded. Insets show magnified views of representative membrane regions, with cluster boundaries highlighted and the number of localizations within each cluster indicated. Rapamycin stimulation markedly increased both the number and density of nanoscale FGFR1 clusters. **(c)** Example of automated cluster selection based on spatial localization density within the reconstructed STORM point cloud. **(d)** Identified individual clusters visualized as color-coded point clouds, each color representing a single detected cluster. **(e)** Quantification of the number of FGFR1 clusters per cell before and after rapamycin stimulation. Rapamycin treatment resulted in a significant increase in cluster number across all analyzed cells. **(f)** Comparison of the number of single-molecule localizations per unit area between background membrane regions and detected FGFR1 clusters. Cluster regions contain substantially higher localization density, confirming successful nanoscale enrichment and validating the cluster identification pipeline.

**Figure S8.**
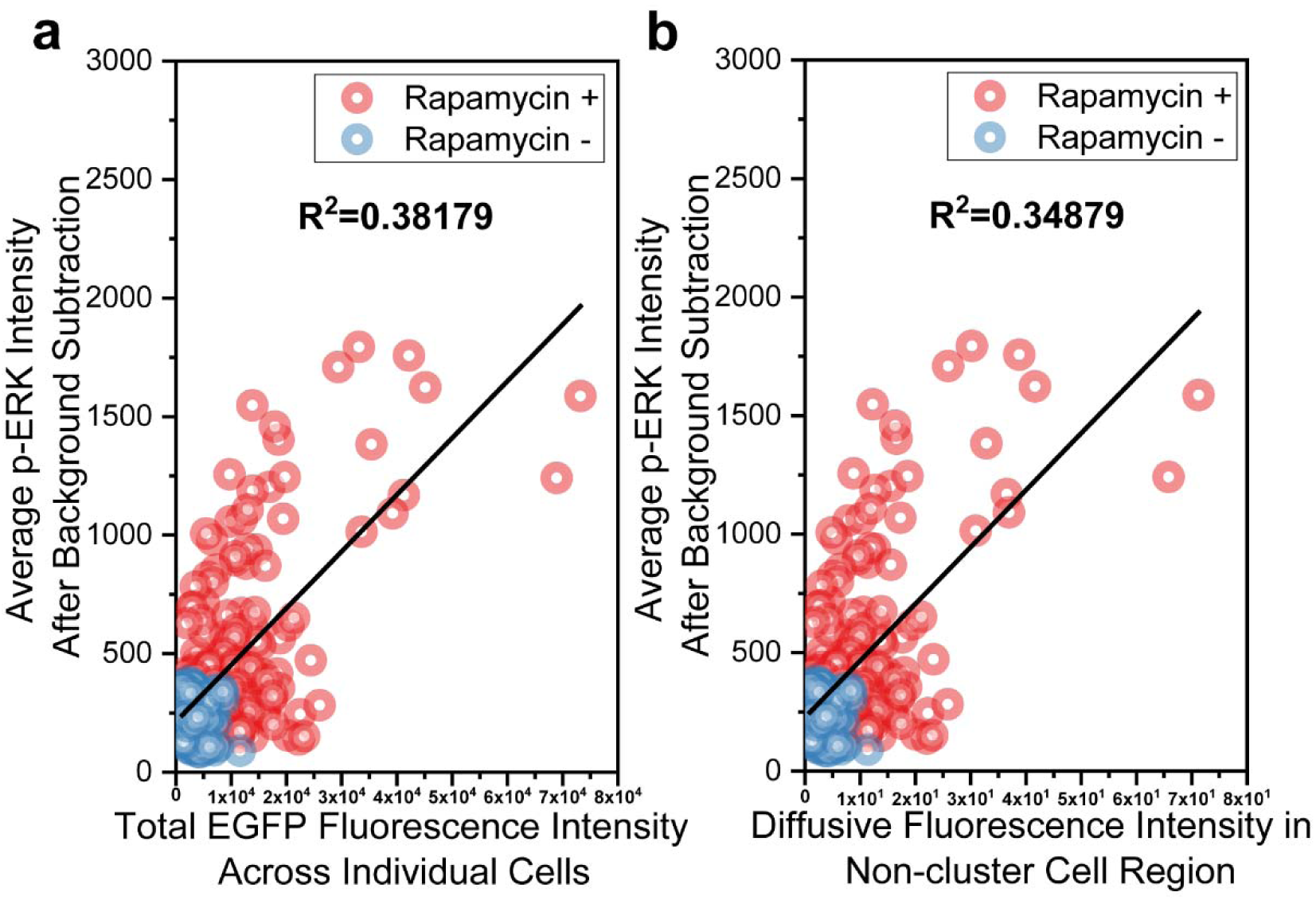
Single-cell correlation of FGFR1 measures with p-ERK **(a)** Correlation between total EGFP fluorescence intensity (i.e., the total FGFR1 amount in each cell) and total p-ERK fluorescence intensity across individual cells. (**b)** Correlation between the diffusive fluorescence intensity in non-cluster cell region (i.e., the FGFR1 amount outside FGFR1 clusters the in each cell) and total p-ERK fluorescence intensity across individual cells. Linear regression analysis shows these correlations are markedly lower than the correlation between FGFR1 amount within FGFR1 clusters and total p-ERK fluorescence intensity across individual cells, indicating that p-ERK activation is primarily driven by FGFR1 clusters rather than by diffusive, unclustered FGFR1.

**Figure S9.**
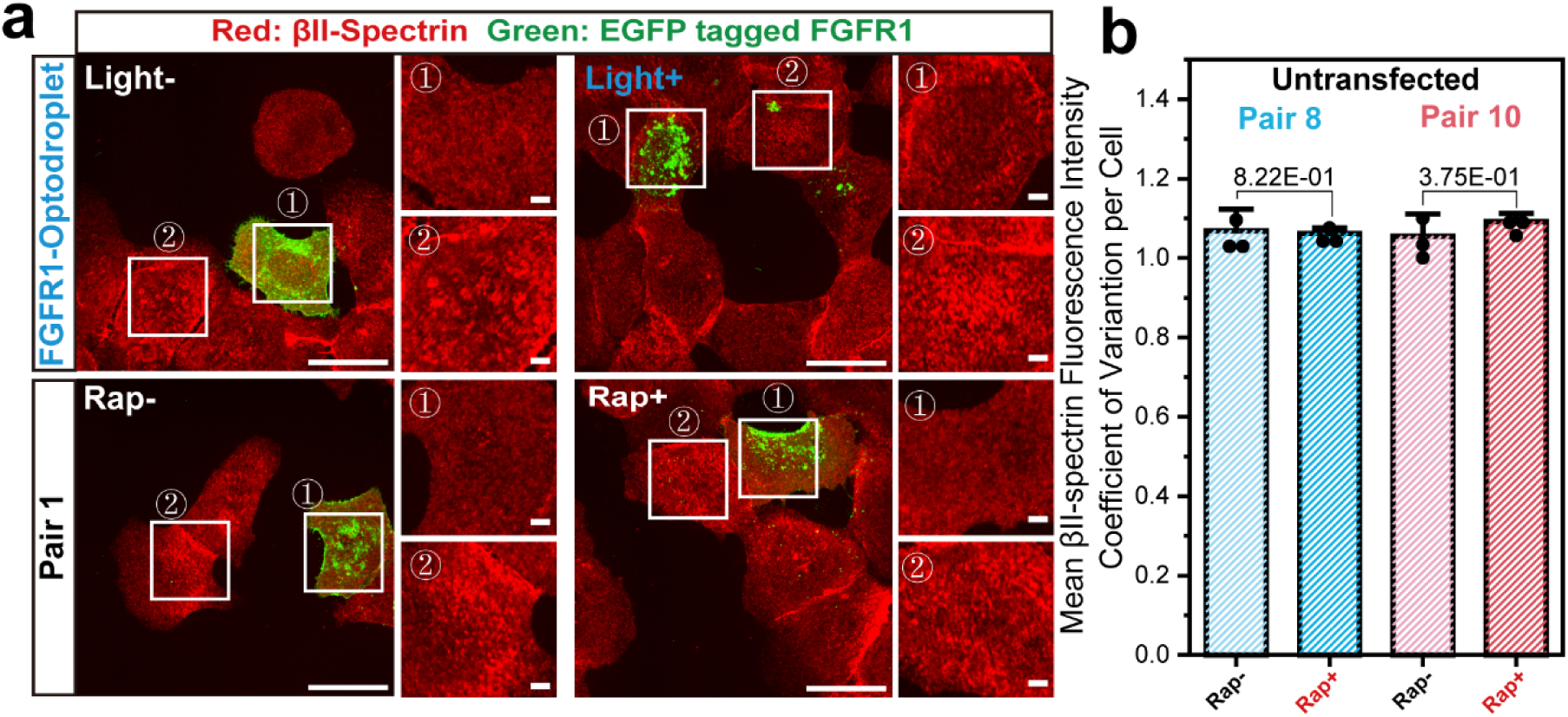
Spectrin degradation in cells expressing optogenetic and chemically inducible FGFR1 constructs with considerable basal ERK activation **(a)** Representative immunofluorescence images of U2OS cells expressing either FGFR1-Optodroplet (top row) or rapamycin-inducible FGFR1 Pair 1 (bottom row), shown under control and stimulated conditions. Cells were stained for endogenous βII-spectrin (red) to assess cytoskeletal integrity. White boxed regions highlight transfected ① and adjacent untransfected ② cells from the same field of view. Optogenetic constructs were stimulated with blue light for 10□minutes, and rapamycin-treated cells were incubated with 500□nM rapamycin for 30□minutes prior to fixation. In both groups, spectrin signal was substantially reduced in transfected cells following activation, whereas neighboring untransfected cells retained intact spectrin architecture. Scale bars, 10□μm (Original view), 2 μm (Cropped view). (**b)** mean coefficient of variation (CV) of βII-spectrin fluorescence intensity for untransfected cells, used as a proxy for spatial heterogeneity and cytoskeletal patterning. Cells with intact spectrin networks exhibit high CVs due to alternating bright/dark regions, and spectrin-disrupted cells show reduced CV. Values are mean ± s.d. from three biological replicates. p-values are calculated using a two-sided unpaired Student’s T-test.

**Figure S10.**
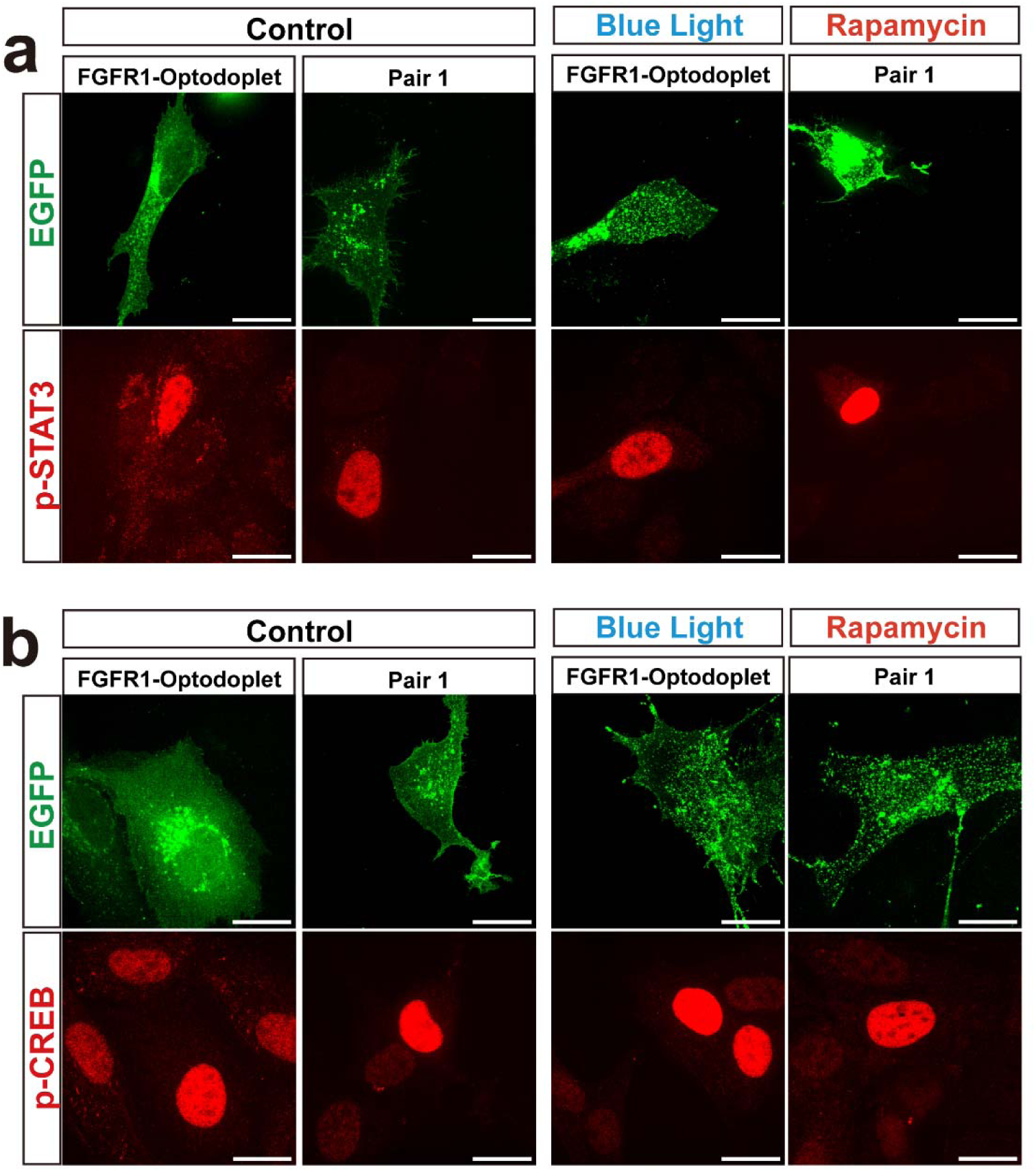
Comparison of downstream STAT3 and CREB activation between optogenetic FGFR1 and chemically inducible Pair 1 constructs. **(a)** Representative confocal images showing p-STAT3 levels in U2OS cells expressing either FGFR1-Optodroplet or Pair 1 under three conditions: untreated control, blue-light stimulation (10 min), and rapamycin treatment (500 nM, 30 min). EGFP fluorescence marks the expression and localization of each construct. In the optogenetic system, substantial background p-STAT3 activation was observed even without illumination and remained elevated upon blue-light stimulation. In contrast, Pair 1 exhibited minimal basal p-STAT3 and showed strong nuclear p-STAT3 induction only after rapamycin stimulation. **(b)** Corresponding p-CREB staining under the same conditions. FGFR1-Optodroplet displayed strong baseline p-CREB activation in the absence of light, whereas Pair 1 maintained low background levels. Rapamycin treatment selectively induced robust p-CREB activation in cells expressing Pair 1, demonstrating the low basal activity and high on-demand responsiveness of the chemical-inducible design compared with the optogenetic system.

## Notes

### Competing Interest Statement

The authors have declared no competing interest.

### Summary of Updates

Figures 1, 2, 3, 5 revised; Supplementary information file updated.

